# The Conductome: A Bayesian Classifier Approach to Predicting and Understanding Behaviour

**DOI:** 10.64898/2026.06.05.730500

**Authors:** Santiago Herce Castañon, Christopher R. Stephens

## Abstract

Predicting and understanding behaviour is a primary objective of many disciplines, especially human behaviour, as it is the cause of many of the world’s most pressing problems. Although it is a fundamental concept in multiple disciplines, there is no agreed operational definition of what it is. Neither is there a generally agreed theoretical framework for predicting it. Here we propose a data-driven approach, using the “Conductome” — the complete set of factors that both predict and explain a behaviour — to operationalise a discipline-neutral definition of behaviour that is based on an ensemble of stimulus/response measurements of a system, showing that it must be determined through a process of statistical inference. As the prediction of behaviour can be characterised as a classification problem, we argue that Bayesian classifiers offer a promising framework in which explainable prediction models that can approximate the Conductome can be developed. We show the efficacy of the framework using a dataset of 1075 persons, with over 3000 features, constructing a model for predicting sedentariness, a behaviour that is a known risk factor for obesity and metabolic disease. We analyse the effect size, coverage, statistical significance and potential causality of a subset of 396 features associated with 58 variables.of different types.

## 1 Introduction

Behaviour is a central concept in both the life and social sciences, including biology, psychology, economics, sociology, anthropology, medicine and health. Indeed, some disciplines define themselves in terms of behaviour: “Psychology is the study of the mind and behavior” [1]; sociology is “the study of social life, social change, and the social causes and consequences of human behavior.” [2]. However, each discipline has its own definitions, as well as a lack of consensus within each discipline, mainly linked to the particularities of the discipline, both in terms of their objects of study - What is behaving? - animals or individual humans, or groups of humans? - as well as the context of the behaviour: economic, versus social, versus health for example. However, besides a plethora of different definitions among disciplines there is also broad disagreement within disciplines. For example, in the case of biology [3], Levitis et al. “found in excess of 25 operationally distinct definitions of the word behaviour” among 181 behavioural biologists! Similarly, in the case of psychology [4, 5], Bergner has emphasised: “Psychology, although describing itself as “the science of behavior,” has not to date arrived at any consensus in the matter of what the concept of “behavior” means.” [4]. Unfortunately, behaviour is one of those concepts that is sufficiently common and intuitive, and with such broad application, that its lack of definition is perhaps less intellectually discomforting than it should be. As in many areas, an attitude of “I know it when I see it” may be deemed sufficient. In the context of human behaviour it is also one of the principal causes of the world’s major problems, such as climate change, environmental degradation, biodiversity loss, emerging diseases, obesity, chronic-degenerative disease, poverty, inequality, social injustice and violence, with behaviour change to ameliorate the adverse consequences being of vital importance. It is also the principal characteristic that distinguishes the physical from the biological.

More important, however, than answering: What is behaviour? is the answer to the question: “Why” is a behaviour? and, in the case that it is a behaviour with adverse consequences, how to change it. Behind the “whys” lies a set of related questions: “Who” is more likely to exhibit a behaviour? “Where” is that behaviour most likely to be observed? “When” is it more likely to be observed? “What” are the stimuli most likely to lead to a behaviour being observed? The word “likely” here is used pointedly, as we will show that both the characterisation of a behaviour and its prediction must be probabilistic, based on an ensemble of measurable events. Distinct to the practice of simply observing behaviour, once it is suitably defined, understanding the “whys” to predict it requires determining the causes, both proximate and distal, that conduce a given organism, as a subject of a certain behaviour in a particular environment, to exhibit it, and a corresponding theoretical model.

A necessary component of a behaviour/conduct is that it links a stimulus and a response, though the stimulus might not be observed, or be unobservable. As such, stimulus-response events can be considered as elements of a class, *C*. Predicting behaviour then becomes a classification problem, where the goal is to predict the probability, *P* (*C*|**X**), of class membership, given a set of conditioning factors, **X**. For a specific behaviour, the question is: Which factors, *X*_*i*_ ∈ **X**, are predictive? with a further question being: Do those factors and resulting theory, or model, explain the behaviour? The universality of the model is then linked to whether those factors can also predict and explain another behaviour and, if that is so - What is the set of behaviours that the model can be successfully applied to? There is no shortage of theories for predicting behaviour and behaviour change. For instance, in a survey in [6] on behaviour and behaviour change across the social sciences, 82 different theories were identified in the literature, with a large part of the diversity being associated with different choices of theoretical constructs on which the theory is based. Of course, it is the power to both predict and explain behaviour across multiple situations and disciplines that must be used to judge the utility of a theory.

In the realm of human behaviour it is clear that there are myriad factors that influence it, ranging from the “micro” of genetics, epigenetics and cell physiology to the “macro” of politics, economics, and other social factors, with all of them adapting over time. This high degree of multifactoriality/multicausality, exhibited over multiple spatio-temporal scales, is the principal manifestation of the complexity of biological systems. It is also what presents the greatest challenges to predicting and explaining them, both from the perspective of obtaining and integrating datasets that exhibit this complexity, as well as having adequate tools for modelling them that capture and respect it. Just as the construction of large bioinformatic datasets has led to the concept of an “-ome” and “omics” [7], representing the idea of a complete set of biomolecules, the obtention of data that represents the complete set of factors that influence a conduct leads us to the concept of a “conductome” [8, 9], where we use the word conduct as opposed to behaviour to distinguish between a purely observational characterisation of behaviour and an understanding of why it occurs, as the etymology of the latter links to the reasons why a certain behaviour is exhibited. ^1^ For the majority of behaviours the associated “why” is often, implicitly, if not explicitly, evolution, both biological and/or sociocultural.

Although one can imagine a complete set of factors from which “all” behaviours may be derived, by completeness in the present work we mean the complete set of factors, **X**_*C*_, that best predict a particular behaviour, where **X**_*C*_ can be different for different *C* and a function of both the subjects exhibiting the behaviour and their environment. In the Conductome the “completeness” of **X**_*C*_ is evaluated via the performance of the prediction model, *P* (*C*|**X**_*C*_), both in terms of predictability and explainability. Another important difference with other omes, both “micro-omes”, such as the genome and metabolome, and “macro-omes”, such as the exposome [10], that represents the totality of an individual’s non-genetic environmental exposures from conception to death, is that in these cases “completeness” is restricted to factors of a certain type, and/or spatio-temporal scale, which, in its turn, is associated with the disciplinary bias inherent in the choice of these factors. In distinction, the Conductome transcends such restrictions by incorporating all factors, without regard to scale or discipline, with a key challenge being to obtain datasets that allow this.

In this paper the principal new contribution is to use the concept and framework of the Conductome to address the questions of: What is a behaviour? and “Why” is a behaviour? With respect to the former, we will argue that the large number of definitions that exist, and the even larger number of theories for it, are a consequence of the wide spectrum of mental models that different practitioners in different fields apply to it. In Bayesian language, everyone has their own Bayesian prior. To avoid disciplinary biases we will develop an operational characterisation of behaviour that is based on observation alone, and applicable across multiple disciplines. In so doing we will also show that it is the principal characteristic that distinguishes the physical from the biological, being an essential property of Complex Adaptive Systems. We will then present a framework, based on the use of Bayesian classifiers and corresponding approximations, for predicting and explaining behaviour that is also applicable across multiple disciplines. This framework is then used to construct a classifier model, *P* (*C*|**X**), to predict a particular conduct — *C* = physical exercise — of particular relevance in health-based applications, using a highly multifactorial dataset with more than 3000 features. We will show that there are many statistically significant factors, associated with different scales and disciplines, that predict the conduct, thereby highlighting the urgent need to develop multiscale, multidisciplinary datasets and the machine learning tools necessary to construct corresponding multifactorial, multiscale, multidisciplinary models.

The structure of the paper is as follows: in section 2 we will address the question of: What is behaviour? beginning with a brief discussion of its characterisation in psychology and biology, then introducing the principal formal elements of our framework: i) defining a statistical ensemble of stimulus-response (cause-effect) events; ii) a characterisation of the distinct roles that the update rules that govern the state changes of a system in the presence of a stimulus play in physical versus biological systems; iii) behaviour seen as a problem in statistical inference; iv) the specification of a suitable benchmark with respect to which a behaviour is defined; iv) statistical diagnostics that quantify differences between stimulus and response in a system versus a suitable benchmark. These elements together lead to a definition of behaviour that is completely operational, as shown in section 2.6. In section 3 we discuss some of the challenges of predicting and explaining conduct, while in section 4 we show that Bayesian classifiers, *P* (*C*|**X**), offer a promising framework within which concrete representations of the Conductome for a given conduct *C* may be constructed thereby allowing for the prediction and explanation of behaviour across multiple disciplines. In section 5 we show how *P* (*C*|**X**) may be computed in high-dimensional situations using machine learning techniques and In section 5.1 we illustrate the feasibility and utility of the approach with a specific health-related example of a conduct — voluntary exercise. Finally, in section 6 we draw some brief conclusions. There is a Supplementary Material 7, where we consider in detail how our approach can help resolve the problem of adequately defining behaviour by applying it to the 20 challenging examples found in [3].

## 2 What is behaviour?

In psychology, behaviour has been defined differently by different scientific organisations. For example, the American Psychological Association provides the following disciplinary definition [1]: “behaviour is an organism’s activities in response to external or internal stimuli, including objectively observable activities, introspectively observable activities and non-conscious processes.” On the other hand, the Oxford Dictionary of Psychology defines behaviour as “the physical activity of an organism, including overt bodily movements and internal glandular and other physiological processes” [11]. In biology, there is no agreed definition and, indeed, disagreement as to whether even a specific biological phenomenon should be considered as a behaviour or not. This has been highlighted in the approach taken by Levitis et al. [3], who presented a list of 20 biological phenomena to 181 behavioural biologists and asked them to characterise them as being behaviours or not, along with a set of statements that represented distinct facets of what a behaviour could be defined to be (see the Supplementary Material S1 for details). They distilled the varied responses into the following definition: “Behaviour is: the internally coordinated responses (actions or inactions) of whole living organisms (individuals or groups) to internal and/or external stimuli, excluding responses more easily understood as developmental changes.” Similarly, Davis et al. [6] used an advisory group of 24 UK experts from the social and behavioural sciences to develop a consensus definition of behaviour. Of course, no one has convened a group of biologists, psychologists, sociologists and economists to form a consensus as to what a behaviour is.

If we ask what is common between the distinct definitions of behaviour from different disciplines, it is the idea that a behaviour is associated with a response of a system due to the action of some stimulus. However, we are then led to ask about the nature of the systems considered: which systems are considered to “behave” and which not? as well as the characterisation of both responses and stimuli. What is clear, however, is that the more detailed nuances of how behaviour is characterised are dependent on both the area and degree of expertise of the experts who define it. Put another way, each discipline and each person is characterising behaviour using a different mental model (Bayesian prior). Below, we will try to characterise behaviour in a way that strips away such prior (disciplinary) knowledge, and is formulated in a mathematical and computational framework that can be made directly operational in terms of observations and data only. The need to operationalise a characterisation of behaviour is also clear in that, from the myriad distinct scientific characterisations of behaviour, even if we were to all accept one, such as the APA definition, we are still left with the problem of how to operationalise it. For instance, just what constitutes an organism’s “activities”? How do we measure them? By organism do we mean plants and animals or just animals? What happens if the stimuli are not observed? In this paper, in the light of how behaviour has been defined in different disciplines, we will develop its characterisation from first principles, rather than from a set of examples and a subsequent consensus of experts. As such, it will be applicable to multiple disciplines. The characterisation is operational in that it depends only on a statistical ensemble of events and the mathematical manipulation of the resulting data. We will show that the characterisation of a behaviour in a system is a problem in statistical inference, where those inferences must be compared to an appropriate benchmark system viewed as a null hypothesis, thus emphasising that behaviour is a “relative” not “absolute” concept. The statistical inference problem is to identify class membership, where the class represents those events relating a stimulus and response (cause-effect) as candidate events for representing a behaviour.

### 2.1 Of Systems, their States and their Dynamics

As the concept of behaviour across disciplines is based on a system responding to stimuli acting on subjects in a given environment, we will first define these concepts operationally. We consider a system, *G*(*t*) = (*S*(*t*), *E*(*t*)), at time *t*, to be composed of a subject, *S*(*t*), and an environment, *E*(*t*), and described in terms of a set of state variables, **X**(*G, t*), which can be separated into subject variables, 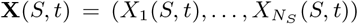, and environment variables, 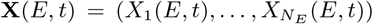. We will initially not restrict behaviour to be only associated with subjects as individuals, but will also consider subparts of individuals as well as groups of individuals. If we define the subject and environment states using very many state variables then we can think of this as yielding a more detailed “microstate” description of the system, and a more coarse grained “macrostate” description when we use only a small number of descriptors. Given a state, *G*(*t*), of the system, we can measure changes of state: Δ_*G*_(**X**(*G, t*^′^), **X**(*G, t*)) : **X**(*G, t*) → **X**^′^(*G, t*^′^), where one or more variables in the subject and/or environment take different values. In order to explain and predict such changes, we can hypothesise the presence of a stimulus, *R*(*t*), specified using a set of state variables, 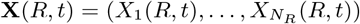, where the variable could be as simple as *R*(*t*) = 0, 1, indicating presence or absence of the stimulus. The state variables of *R*(*t*) can also be considered as part of the subject and/or environment variables, but we will here separate them out. If we wish to posit that *R*(*t*) is the cause of the change Δ_*G*_(*t*^′^) we must have *t < t*^′^ to maintain causality. A state of the world, *W* (*t*), that accounts for the states of *S, E* and *R*, is *W* (*t*) = (*S*(*t*), *E*(*t*), *R*(*t*)). We can define Δ_*G*_ as the response to the stimulus. As a result of the interaction between *R* and Δ it may also occur that *R* itself changes, i.e., there is a change in one or more of the stimulus variables.

We assume that the potentially varied and complex set of processes by which a stimulus is converted into a state change can be modelled by update rules: *U*_*iα*_ : **X**(*W*_*α*_, *t*) → **X**_*i*_(*W*_*α*_, *t*^′^), where *α* = *S, E, R* and the index *i* is to indicate that there are potentially multiple update rules, even in the presence of the same stimulus. Indeed, we will argue below that this is a defining difference between physical and biological systems. *U*_*iα*_ is a representation of the potentially varied and complex set of processes by which a stimulus, as cause, is converted into a state change, as effect, in any of the three components of the world. The state variables themselves may be explicitly measurable — ordinal or categorical, such as position, temperature, gender, weight, income, etc., in the case of subject variables, or they may also be latent variables, such as depression, anxiety, happiness, anger, or other emotional constructs, as measured by corresponding psychological instruments. The stimulus can also lead to changes in environmental variables, such as the presence of a spider’s web, a bird’s nest or more sales of soda in a particular fast-food chain. *U*_*iα*_ is also where the natural representation of mental processes lies, as well as physiological ones.

To illustrate the important differences in the concept of behaviour between physical and biological systems, consider dropping two balls of different mass from a certain height, describing their state by the coordinates and velocity of their centres of gravity, **X**(*S*_*i*_, *t*) = (*x*_*i*_(*t*), *y*_*i*_(*t*), *z*_*i*_(*t*)) and (*v*_*i*_(*t*), *v*_*i*_(*t*), *v*_*i*_(*t*)), and ignoring other details. The stimulus in this case is gravity and the corresponding update rule is Newton’s Second Law. If we know the initial conditions, (*x*_*i*_(0), *y*_*i*_(0), *z*_*i*_(0)) and (*v*_*i*_(0), *v*_*i*_(0), *v*_*i*_(0)), of each ball, we can describe each subsequent state change, i.e., the change in the position of their centre of gravity, determining that the two balls “behave” in the same way, for example, as determined by both falling to the ground and touching it at the same time. If a third object, a cat, is dropped and its state is described only by its center of gravity, it follows the same physical trajectory as the balls. i.e., it exhibits the same “behaviour”. However, if the state description includes the position of its legs, say *X*_*l*_(*t*) = *up, down*, we observe that the cat alters its leg position to land feet first, demonstrating a biological update rule distinct from the universal physical laws governing the balls. Hence, how we characterise a behaviour, through responses modelled as state changes, depends on the set of state variables chosen. Note that, in this example, the state of the environment **X**(*E, t*) is effectively constant. Of course, a complete, microscopic description of the state of a subject and/or environment, **X**_*T*_ (*t*), is not necessary when trying to associate state changes with a given behaviour. However, it is important to identify those variables (degrees of freedom) which are relevant. A description of the three falling objects in terms of their centre of gravity is not sufficient to distinguish them — their “behaviour” is the same as all are slaves to the laws of physics. However, the cat possesses other state variables, and corresponding update rules, that allow us to say it has a different behaviour than the balls, a behaviour that is equally in accord with physical law as the act of falling itself. However, the role of physical law in both cases is quite different. In the case of the centres of gravity, Newton’s Second Law not only predicts the state changes of all three systems but also explains them. However, in the case of the cat, Newton’s Second Law neither predicts nor explains the complex physiological processes that led to the result and, much less, why the cat does it. The force of gravity is both a proximate and ultimate cause of the changes in the centre of gravity of the ball and cat. It is neither a proximate nor ultimate cause of the cat’s righting reflex.

Although there exist sophisticated physics models of deformable bodies that allow us to understand how the fact that a cat can land feet down is consistent with the laws of physics [12], and could be viewed as a proximate cause of the cat’s righting reflex, they do not describe the complex physiological processes that are the true proximate causes that led to the result and, much less, the ultimate cause of why the cat does it, which is the effect of Natural Selection.

It is important to note that, more often than not, the update rule for a given behaviour is not manifest, but must be inferred from its associated state changes. For instance, the explicit rule for updating the states of falling objects was unknown until Newton came along. Of course, this did not prevent people from being aware of the regularities in the motion of falling objects, and being capable of predicting them — people were catching falling apples long before Newton. Indeed, before Newton, the true stimulus — gravity — that caused bodies to fall was not known, but, rather, the Aristotelian nature of “earth” and “water” elements was invoked. The update rule associated with Newton’s Second Law is very simple. However, the update rule that governs the contortions of a falling cat, as the net result of a very large set of complex physiological processes, is not known explicitly. Indeed, in most human or animal activity the update rules associated with state changes are not explicitly known either.

### 2.2 The nature of “choice” in physical versus biological systems

The laws of physics and chemistry such as Newton’s Second law, Maxwell’s equations or the Schrödinger equation, serve as update rules that are universal across the classes of systems to which they are applicable, although the relevant state variables can be quite different among different systems. For instance, coordinates of the centre of gravity and Euler angles, in the case of rigid bodies, are quite different from velocity, pressure and density in the case of Newtonian fluids. However, in both cases the relevant update rule is Newton’s Second Law. Universality in this case refers to the fact that for rigid bodies and Newtonian fluids there is no other update rule than Newton’s Second Law. In biological systems, however, there exist multiple update rules associated with different stimuli, and different update rules for the same stimulus. For example, hunger can cause a state change by making an organism seek food. However, the final states are myriad, depending on what was eaten, how much, where and when etc. It can also depend on the strength of the stimulus, i.e., how hungry is the organism? Hunger itself can also be viewed as a complex change in physiological state (hormonal and neuronal) with a corresponding update rule. However, the relation between stimulus (hunger) and response (e.g., eating a doughnut) is very different to that for the centre of gravity of the falling balls, or the falling cat, where the stimulus is gravity and the unavoidable response is — fall at 9.8*ms*^2^.

The principle difference is that balls and cats, have no “choice” but to fall. However, an organism can “choose” to forage or not, or to eat one food versus another, or to eat one quantity versus another, where by “choice” we do not mean freewill per se, but, rather, to the fact that there exists more than one response to the stimulus and therefore more than one update rule, with each rule linked to a particular state change. Just as the stimulus hunger can lead to a choice among multiple responses, so can multiple stimuli lead to the same response. For example, for smoking or eating events, as advertisers and marketers well know, multiple sensory triggers can lead to purchase of a product. One might think that in these terms a flipped coin or a rolled dice has “choices”, as flipping the coin — the “same” stimulus can lead to multiple responses — heads versus tails. Similarly, we can see that multiple stimuli, flipping the coin in different ways, can lead to the same response — heads. In these cases however, the appearance of choice is illusory. Only Newton’s Laws offer the appropriate update rule, although the actual stimuli — what were the exact forces exerted by the fingers while flipping the coin and when it landed on the floor — are sufficiently complicated that a deterministic prediction of the outcome is infeasible. Of course, an ardent proponent of determinism might argue that the biological cases are just much more complicated versions of flipping coins. However, in physical systems we know the underlying update rules, and the relation between stimulus and response is clear and direct, just difficult to measure. Although we might imagine that Newton’s laws, or the Schrödinger equation, could offer a universal update rule for biological systems, in terms of making predictions and explaining their complex, adaptive behaviours, they are of very little to no use.

To distinguish behaviour in living systems from physical systems we will now restrict the term behaviour to be only associated with systems that have multiple update rules, and therefore different behaviours, in the presence of the same stimulus. Thus, behaviour is characterised by a state change in a subject and/or environment that is brought about by an update rule: *U*_*iα*_ ∈ **U**, *U*_*iα*_ : **X**(*W*_*α*_, *t*) → **X**_*i*_(*W*_*α*_, *t*), where **U** = (*U*_0_, *U*_1_, …, *U*_*m*_) is the set of possible update rules for *S* and/or *E*. Within these update rules is a rule, *U*_0_, such that there is no state change with respect to the considered state variables. An intuitive example of how a given state change can result from many different update rules is financial trading, where a given state change, for example — buying 1000 shares of a given stock — can be a result of many different update (trading) rules. A rule may be algorithmically explicit, such as “buy when price is lower than a fixed threshold”, or may be more complex, based on a sophisticated mental model associated with experience and intuition. In this sense, there is no one-to-one map between update rule and state change. Different trading rules can lead to the same response — buy 1000 shares of a given stock — and therefore with respect to that single event are not distinguishable. However, that does not mean that applied multiple times they lead to the same results. Similarly, when hungry, a person may choose to eat a doughnut, eat a piece of fruit, or eat nothing at all. Thus, in contradistinction to physical systems that have states and unique update rules that describe the transition from one state to another, biological systems are associated with multiple update rules and their dynamics takes place within a space of states and rules, not just a space of states^2^ Given our emphasis on the existence of multiple update rules, and therefore responses, in the presence of the same stimulus it might be wondered how can falling cats be considered to have a “choice”? The answer to this question will be given in the following sections, where we will argue that a behaviour is a “relative” not ”absolute” concept and must be quantified and interpreted in the context of a statistical ensemble of events.

### 2.3 Identifying Behaviour is a Statistical Inference Problem

All the examples of behaviour of [3], as shown in S1, refer to one specific event that associates the response of a subject to either an explicit or implicit stimulus. However, if we strip away any prior knowledge of the phenomena, how do we know that the relation between supposed stimulus and supposed response is not just a random occurrence? What degree of correlation between stimulus and response is required to determine that there is a relation? For example, imagine, from a zero prior knowledge state, one observes a cat falling from a tree, twisting and landing on its feet. Would this be sufficient evidence to assert a behaviour that characterises falling cats as always landing on their feet? If we observed 100 different cats falling from trees, and they all landed feet first, would that be sufficient evidence? What about 60? or, we observed the same cat, unfortunately, falling 100 times from the same or different trees? What about an ensemble of 50 cats and 50 dogs? Again, to answer these questions we must first try to eliminate all prior knowledge of falling cats, or dogs, and try to characterise their different behaviour from pure observation. Thus, establishing a statistically significant correlation between stimulus and response is a necessary condition for potentially calling their relation a behaviour, which means that characterising behaviour is a problem in statistical inference, and must be addressed in the framework of a statistical *ensemble* of events, 𝒱. Of course, there is a vast literature in ethology and psychology, for example, where specific behaviours in specific systems have been observed and quantified using statistical analysis of experimental data. However, this is not the same as using a probabilistic framework in order to define and predict behaviour generally.

An ensemble of events that is generated through the design of experimental protocols designing we will call an “external” ensemble, 𝒱^*e*^. We claim that the existence of ensembles is an intrinsic property of our intuition about behaviour, but is implicit rather than explicit, being based on an “internal” ensemble of events, 𝒱^*i*^. Thus, typical assertions about behaviours, such as:“Frank exercises regularly”, “Joanne eats unhealthily”,“James buys tech stocks”, “cats land feet first”, have implicitly associated with them the notion of an ensemble of “observations”, as well as a corresponding null hypothesis with respect to which the probability of the behaviour is judged. On meeting Frank or Joanne for the first time, while Frank was jogging or Joanne was drinking a large, high-calorie soda, we would not be likely to make assertions about Frank or Joanne’s behaviour based on one observation. However, if we had known Frank and Joanne for a long time, and had observed Frank going to the gym five times a week, or had observed Joanne drinking several bottles of high-calorie soda every day, then we would be much more likely to make such assertions. However, in making these judgements there is always in the background a baseline. For example, if Frank exercised on 12 random days in the year we would not be likely to describe Frank’s behaviour as “exercising regularly”, while if he exercised on 250 days in the year, random or not, we would be more likely to make the assertion. Similarly, if Frank exercised 52 times a year, but always on Monday, we would be more inclined to call it a behaviour than if it was on 52 random dates. In these latter two cases we might also be inclined to call the behaviour a habit or routine. Note that in none of these cases have we introduced an explicit stimulus for the responses. However, in our mental models we often posit potential stimuli from previous knowledge, such as: “Frank exercises because he wants to keep fit”; whereas, if Frank was overweight, we might posit “Frank exercises because he wants to lose weight”. Moreover, even for a single observation, we must specify conditions that define what we mean by a co-occurrence of stimulus and response. For example, it may be that the response, Δ(*t*^′^), to a stimulus, *R*(*t*), is such that it occurs a substantial amount of time after the putative stimulus has been observed. How, then, do we know that the response was due to that stimulus? It may be that the stimulus is an indirect cause, as opposed to a direct one. Worse, if there were other candidate stimuli that occurred in the time interval between *t* and *t*^′^, how do we determine if one of these other stimuli was responsible for the response rather than *R*(*t*)?

More formally: for a given subject, *S*, and environment *E*, we consider a statistical ensemble, 𝒱, of events, *v*_*C*_ (*t*) ∈ 𝒱, where we introduce now a class variable, *C*, to represent the co-occurrence, (*R*, Δ), of a stimulus (cause), *R*(*t*), and a corresponding response (effect), Δ(*t*), or, in the absence of an explicit stimulus, the occurrence of a state change Δ(*t*^′^). The state change Δ(**X**(*W*_*α*_, *t*^′^), **X**(*W*_*α*_, *t*)), can occur in the subject, the environment, or both, via an update rule: *U*_*iα*_ : **X**(*W*_*α*_, *t*) → **X**(*W*_*α*_, *t*^′^). Both *R* = *R*(*S, E*) and Δ = Δ(*S, E*) are functions of the subject and/or environment considered. Indeed, as mentioned, the stimulus *R* can also be represented as a subset of the subject and/or environmental variables. In the former case, *R* could represent hunger, while in the latter it could be the presence of nutrients. As mentioned, it may be that the stimulus and update rules are not explicitly present, or measurable. It may also occur that the stimulus has a direct impact on a set of state variables that are not directly measurable and that the link between stimulus and state change is indirect and inferential. We can construct a statistical ensemble, 𝒱, in different ways: i) a longitudinal ensemble, where a series of observations of events associated with the same subject is made over time; ii) a transverse ensemble, where a series of observations of events associated with a set of subjects, and/or environments, is made; iii) a mixed ensemble, where a series of observations of events associated with a set of subjects and/or environments is made over time. Just as the characterisation of a behaviour is dependent on the state variables used to describe the system, so it also depends on the ensemble of events used, as the statistical characterisation of the relation between stimulus and response depends both on the number of observations and on the type of events that are included. Note that in constructing this ensemble, although we can imagine a set of explicit, empirical observations of the events themselves, the corresponding state changes in the subject and/or environment and, potentially, the corresponding update rule and the stimulus that led to the state change, the intuition, and resulting ambiguity, of behaviour, as relating to certain relations between stimuli and responses, is associated with an “ensemble of observations” that is a mental construct. Such mental constructs are what we use to characterise and distinguish behaviours. We will refer to ensembles based on empirical data as external, or extrinsic, ensembles, 𝒱^*e*^, while those that are inherent in our mental constructs of phenomena as internal, or intrinsic, ensembles, 𝒱^*i*^.

If for a subject, *S*, and environment, *E*, we identify from the ensemble, 𝒱(*S, E*), of *N* events, a subset, 𝒱_*C*_ ⊂ 𝒱, of *N* (*C*) events, where a particular state change, Δ(**X**_*C*_ (*t*^′^), **X**_*C*_ (*t*)), has been observed in the system and/or environment, and where **X**_*C*_ represents a subset of the set of potential descriptors of the state, we may ask how frequent is the event *v*_*C*_ ∈ 𝒱_*C*_ . Although this depends on our choice of 𝒱 in the first place, for *N* events, however defined, if *N* (*C*) is the number of events where the behaviour is observed then the probability for that behaviour is *P* (*C*) = *N* (*C*)*/N* .

### 2.4 Behaviour is Relative

We will now add another condition necessary for characterising behaviour: that it is “relative” not “absolute”, meaning that implicitly, and sometimes explicitly, behaviour, as a probabilistic response to a stimulus in a system (*S, E*) is characterised by comparing the response to some benchmark system (*S*^′^, *E*^′^). Thus, for every class event, *C*, that potentially characterises a behaviour, we must compare *P* (*C*|𝒮,ℰ) with *P* (*C*|𝒮^*t*^, ℰ^*t*^) for some benchmark. Hence, behaviour is intrinsically an attribute that distinguishes one system, or set of systems, (𝒮, ℰ), from another system, or set, (𝒮^*t*^, ℰ^*t*^). This is manifest in those examples in the Supplementary Material, taken from [3], where there is no explicit stimulus. When there is a stimulus we can always compare *P* (Δ|*R* = 1), when the stimulus is present, to *P* (Δ|*R* = 0), when it is absent, or to *P* (Δ), which are natural benchmarks. In this sense, all behaviour is behaviour “change”, or, rather behaviour “difference”, meaning we compare the responses of a system in two different situations, one where the stimulus is absent and another where it is present. However, in the absence of explicit observations of the stimulus we must use a different benchmark. In analogy with presence/absence of the stimulus this could be presence/absence of a certain subject type, or presence/absence of a certain environment type. We can, for example, compare potential behaviours, keeping *E* fixed for system and benchmark, and comparing responses of one subject *S* with another *S*^′^. Similarly, we can keep *S* fixed and compare responses in one environment *E* with another *E*^′^.

The scope and context of a given behaviour determine the appropriate benchmark system against which it must be compared. Thus, flying is a behaviour that distinguishes birds from reptiles but not birds from bats. Falling and landing feet first is a behaviour that distinguishes cats from dogs, but not Siamese cats from Persian cats. It might seem unsatisfactory that we require such an apparently subjective element as a choice of benchmark system in order to characterise a behaviour. However, we would respond by saying that in biology and psychology that subjectivity is already present in our mental models of behaviour, seen as Bayesian priors, and is the reason for the large number of definitions that exist, and for the wide range of responses seen in [3]. Here, we are just bringing to the fore and making explicit the presence of a benchmark. With respect to what is a reasonable benchmark, it should be chosen so that it represents a system that is “close” to the system for which we are trying to characterise its behaviour, where a quantification of “close” requires a metric on the space of systems, a topic we will return to in the future. Here, we will rely on the intuition that is already present in our mental models. For instance, intuition would lead us to posit that falling dogs might be a reasonable benchmark for considering falling cats landing feet first as a behaviour, whereas falling fish or falling birds would not be, with a principle element being associated with the benchmark possessing, at least, the state variables associated with the system under study. Equally, we would not naturally consider falling Persian cats as a reasonable benchmark for a system with Siamese cats as subjects. In S1 we show which of the examples of [3] fall into our characterisation of behaviour with respect to an appropriate benchmark and which do not, emphasising that a stimulus and response can be appropriately viewed as a behaviour with respect to one benchmark but not necessarily with respect to another..

Behaviour as being relative versus absolute is also fundamental to the concept of behaviour change in time. For instance, going from smoking to not smoking. To speak of a change in behaviour, i.e., the its probability goes up or down, requires a benchmark. For instance, a before and after a particular intervention. This relativity of behaviour is strongly linked to its potential interpretation as an adaptation. Thus, a subject can have different behaviour to another as an adaptation, or a subject can have different responses to different environments. Again, as an adaptation. Indeed, we would argue that all behaviour is associated with behaviour change in this sense, with the prime “intervention” being natural selection itself. It is also this that can cause confusion in our mental models of behaviour, as it is more natural to think of a behaviour as such when it has been learned, but less so, in many cases, when it is innate, as in the latter the “choice” element that we have emphasised in the relation between stimulus and response is less clear. Innate behaviours, such as cats falling and landing feet first, apparently challenge our assertion that behaviours only occur in systems with multiple update rules. Does the cat have a “choice” in landing feet first or not? In contrast with starting or stopping smoking, or eating healthily or unhealthily, or playing tennis versus golf, we think of the former as innate behaviour and the latter as learned behaviours.

For innate behaviour *P* (*C*|*S, E*) ∼ 1, as, for example, in the case of falling cats as subjects, *S*, in an environment, *E*, represented by falling from a height. However, if, for the same group of cats, *C*_*i*_ represents as a behaviour their consumption of different cat foods, *i* = 1, …, *m*_*f*_, then we would not find *P* (*C*_*i*_|*S, E*) ∼ 1, but a distribution of probabilities that represents their food preferences. Moreover, by reinforcement learning this latter behaviour of food preference can be changed. i.e., the cat can learn to prefer one food versus another. On the other hand, the reflex of landing feet first cannot. In other words, food preference as a behaviour is more “plastic”, i.e., more amenable to change, than landing feet first. In general, there is a spectrum of degrees of plasticity associated with different behaviours as a function of time, subject and environment, which can be quantified, as we will show in section 2.5. We can first think of plasticity as a function of *S* and *E*, and as such it is dependent on our choice of ensemble 𝒱. As a function of *S* plasticity is associated with the degree of heterogeneity of the probability of the behaviour being observed over different populations. Similar notions apply for plasticity as a function of *E*. The bigger the difference (*P* (*C*|*S, E*) − *P* (*C*|*S*^′^, *E*^′^)) between one system and another the more plastic is the behaviour between the two groups.

However, the concept of behaviour plasticity is even more appropriate in the context of behaviour change in time. In this context we may imagine a specific intervention, *I*, such as a smoke cessation program, viewed now as a stimulus that can lead to a state change, and consider the probability of observing the behaviour in the presence of the intervention versus a benchmark where it is absent, or where it is marginalised. In this case, for example, *S* could represent a given population of subjects, smokers say, that received the intervention, and *S*^′^ those who did not, and where we set *E* = *E*^′^ so they are in the same environment. In this example, we would want *P* (*C*|*I, S, E*) *< P* (*C*|*S*^′^, *E*), i.e., that the intervention reduced the incidence of smoking. Thus, the bigger the difference (*P* (*C*|*I, S, E*) − *P* (*C*|*S*^′^, *E*)), the bigger the impact of the intervention (stimulus). To try and isolate the effect of the intervention, as in a clinical trial, we would also prefer to have all the considered state variables of *S* to be the same as those of *S*^′^ except that in *S* the intervention was applied.

If we try to extrapolate this analysis of behaviour change to a more biological case, the principle difficulty is in identifying an explicit intervention and interpreting it. Natural selection itself can be interpreted as the intervention *par excellence*, either through its causes or its consequences, both direct and indirect. In some circumstances a behavioural change can be traced to a particular genetic mutation [16, 17, 18], although the more complex the organism the more difficult it is to make a distinction between the multiple potential causes of a behaviour difference, as is the case for example with behavioural adaptations in dogs [4]. However, innate behaviours are not eternal, in that there was an evolutionary past, where, before a given epoch, the behaviour was not present, and a later epoch when it was. Thus, for falling cats, we can hypothesise that there were populations of ancestors of cats as subjects, *S*^′^, before they became arboreal, that did not possess the righting reflex. Passing to an arboreal existence led to a selective pressure that had as consequence a preference for those animals that developed the physiological and behavioural adaptations that constitute the righting reflex, which we can take as subjects *S*. Thus, at some point in time *P* (*C*|*S*^′^, *E*) ∼ 0 and later *P* (*C*|*S, E*) ∼ 1. To quantify this we require a longitudinal ensemble that incorporates *S* and *S*^′^ . Clearly, between those two points in time the probability is changing, reflecting the fact that there is a behaviour change. With this in mind, we would argue that the difference between innate and learned behaviours is really a question of the timescale over which behaviour has occurred and the degree of positive and negative feedback to which an individual or population of subjects exhibiting the behaviour has been subject. In other words, innate behaviour itself is also “learned”, but over timescales of, potentially, many generations as distinct to learning within a lifetime of an individual. Additionally, the ensemble within which the behaviour change is observed may contain different species if it is the case that the adaptations were associated with a speciation event.

### 2.5 A Frequentist Perspective on Behaviour and Behaviour Change

For an external ensemble, 𝒱, of *N* events associated with a set of subjects and environments, (*S, E*), a behaviour is associated with a subset 𝒱_*C*_ ⊂ 𝒱 of *N* (*C*) events, where the class *C* is definable in terms of co-occurrences of stimulus and response, (Δ, *R*), with *P* (*C*) = *P* (*R*, Δ) = *N* (*R*, Δ)*/N* = *N* (*C*)*/N* . To determine how the behaviour changes, or is associated with different characteristics of the subject and/or environment, we can consider subsets of events, *N* (**X**(*S, t*)), *N* (**X**(*E, t*)) or *N* (**X**(*S, t*), **X**(*E, t*)), where we restrict to certain subjects or environments by fixing particular subject and/or environmental variables. This is done by incorporating more of the state variables, **X**(*S, t*), and/or **X**(*E, t*), in the characterisation of the class events. For example, we might consider **X**(*S, t*) to just be a binary label that denotes whether *S* is a cat or not. We could then add in, for example, the cat’s age, to see if the behaviour was age dependent. How many, and which, variables should be included is dependent on the questions we wish to ask. In the context of human behaviour there are many variables, both subject and environment, that are potentially of interest. For example, in the case of a behaviour such as consumption of “junk food”, in the context of obesity and metabolic disease, we are interested in many characteristics of the subject — education, age, socio-economic factors, genetics etc., — and the environment — food availability, economic system, public health policies etc. One reason for an interest in all these factors is the identification of possible causes or risk factors for the behaviour and the search for appropriate interventions.

We can also count events, *v*_*C*_, on another sub-ensemble, 𝒱(*S*^′^, *E*^′^), where (*S*^′^, *E*^′^) can represent different subjects and/or environments, and which will be used as a benchmark. For example, out of *N* events for a population of males (*S*) and females (*S*^′^) of a given species in the same environment, for a given stimulus-response, or just response, *C*, we can calculate and compare *P* (*C*|*S, E*) and *P* (*C*|*S*^′^, *E*), which will tell us how likely males are to respond to the stimulus *R* versus females. Note that the ensembles 𝒱(*S, E*) and 𝒱(*S*^′^, *E*^′^) may be longitudinal, transverse or mixed. Indeed, it is precisely the longitudinal and mixed case that are of particular interest in health-related applications, where, for example, the class variable, *C*, is smoking, and *S* represents a group of smokers before an intervention, such as a smoke ban in public places, and *E*^′^ the environment after the ban. (*P* (*C*|*S, E*) − *P* (*C*|*S, E*^′^)) is then a measure of the impact of the smoking ban on the frequency of the behaviour — smoking. Thus, our fundamental object of interest is *P* (*C*|*S, E*), the probability to observe the class events *C* conditioned on the subject and environment in which the events are being observed. If we are to argue that behaviour is associated with comparing a link between stimulus and responses compared with a benchmark, then given the probabilistic nature of the link, this is a statistical inference that must be tested using a suitable diagnostic. An appropriate one is a binomial test, where we take as null hypothesis *P* (Δ)

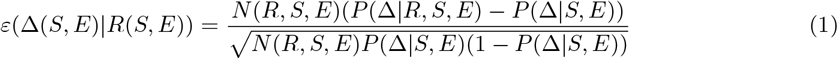

where, for simplicity, on the right hand side we have suppressed the dependence of *R* and Δ on *S* and *E*. Thus, we consider an ensemble, 𝒱_*R*_ of *N* (*R, S, E*) events, where the stimulus *R* is observed, and we count those events, (Δ, *R*), where the response, Δ, is also observed, remembering that these events require an operational definition of the co-occurrence of stimulus and response. The null hypothesis is that the probability of the response is independent of the stimulus. Although this could be posited as a Bayesian prior, it can also be measured if we consider in our ensemble, 𝒱, events, (Δ, 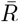) where the stimulus is absent, 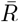, with probability 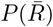. In this case, 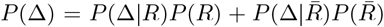. We could, of course, also have chosen as null hypothesis 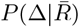— how often the state change is observed in the absence of the stimulus. As the intuition of a behaviour is that it is a predictable response to a stimulus, we might be less inclined to call it a behaviour if (*P* (Δ|*R*) − *P* (Δ)) ∼ 0.

If the stimulus, *R*, is a sufficient condition for generating the response, Δ, in the system (*S, E*), then *P* (Δ|*R, S, E*) = 1. We might want to claim this as definitely a behaviour, as the stimulus always leads to the response. However, this does not imply that it is also necessary. If it is also necessary, then *P* (*R, S, E*|Δ) = 1. The ratio *P* (Δ|*R, S, E*))*/P* (Δ|*S, E*)) is then a measure of the degree of sufficiency of *R*, while *P* (*R, S, E*|Δ))*/P* (*R, S, E*)) is a measure of its necessity. If both *P* (Δ|*R, S, E*) = 1 and *P* (*R, S, E*|Δ) = 1, then the stimulus *R* is both necessary and sufficient to observe the response Δ, at least in the context of the particular subject and environment considered. If we have *P* (Δ|*R*) ∼ 1 and 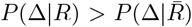 but 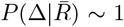 too; in other words, the response is observed both when the stimulus is present and when it is absent, but somewhat more frequently when it is present, then *R* is sufficient but not necessary. This is a very common occurrence in behaviours, indicating that there are other stimuli (causes) that can lead to the same response, Δ. Similarly, it may occur that a condition is necessary, *P* (*R, S, E*|Δ) = 1, but not sufficient, i.e., *P* (Δ|*R, S, E*)) *<* 1, indicating that the same stimulus can lead to multiple responses. As argued, it is the existence of multiple responses to the same stimulus, and multiple stimuli leading to the same response, that mark out biological systems from physical systems.

In equation (1), when the binomial distribution can be approximated by a normal distribution, |*ε*| *>* 1.96 corresponds to the 95% confidence interval that the observed relation between stimulus and response is not consistent with the null hypothesis. Although the value of *ε*(Δ|*R*) is completely objective, its interpretation through the choice of benchmark is not. For example, what, if any, threshold would we impose on the difference (*P* (Δ|*R*) − *P* (Δ)) in order to think of it as a behaviour?

When the stimulus cannot be explicitly observed or measured, and we have only events where there was an observed state change, denoted by the class variable *C*, we can consider two groups of events in any of our external ensemble types. One group will be denoted by the combination (*S, E*) and another with the combination (*S*^′^, *E*^′^), with both combinations specified using the corresponding sets of state variables. We then consider the number of occurrences, *N* (*C, S, E*), of the state change *C* in the set of events denoted (*S, E*), which is a sub-ensemble of 𝒱. We can then compare *P* (*C*|*S, E*) to a benchmark (null hypothesis), *P* (*C*|*S*^′^, *E*^′^). To test the hypothesis that the probability of the behaviour *C* in the subject *S* and environment *E* is statistically significantly different from the probability of observing it in the subject *S*^′^ and environment *E*^′^ we use

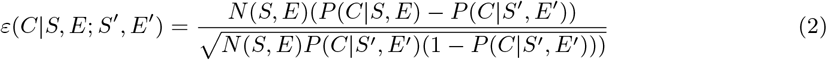

The underlying question that equation (2) answers is: Is the behaviour of a certain subject (subpart, individual or population) in a given environment is significantly different to that of another subject, potentially in a different environment, if *E* =/ *E*^′^? If we choose *S* = *S*^′^ we can analyse differences in behaviour as a function of environment, or choose *E* = *E*^′^ and analyse differences as a function of the subject type. Note that in equation (2) we are now working with an ensemble of events 𝒱(*S, S*^′^, *E, E*^′^) = 𝒱(*S, E*) ⊕ 𝒱(*S*^′^, *E*^′^). It is within this ensemble that the appearance of “choice” arises as, even though on 𝒱(*S, E*) we may have *P* (*C*|*S, E*) = *N* (*C, S, E*)*/N* (*S, E*) = 1, on 𝒱(*S, S*^′^, *E, E*^′^), *P* (*C*|*S, E, S*^′^, *E*^′^) *<* 1.

To illustrate the use of (2), consider *S* to be a heavy ball and *S*^′^ a lighter one that we will let fall under the force of gravity and we also have *E* = *E*^′^. Many different response variables could have been chosen, from: “did it fall = yes/no” — to a measurement of the time from release to hitting the ground, with the different response variables providing different perspectives on the behaviour of the balls in the gravitational field. For instance, the time from release to impact would provide information about their rate of fall when compared to whether the ball fell or not. Our ensemble of events is *N* (*S, E*) repetitions of dropping the heavier ball. We observe *N* (*C, S, E*) events where the response is the same and we compare that to the expected number *N* (*S, E*)*P* (*C* | *S*^′^, *E*^′^) if the null hypothesis were correct. So, how do we calculate *P* (*C* | *S*^′^, *E*^′^)? It can be posited as a Bayesian prior, or it could be measured using an ensemble of *N* (*S*^′^, *E*^′^) events, counting among them the class events *N* (*C, S*^′^, *E*^′^), and calculating *P* (*C*|*S*^′^, *E*^′^) = *N* (*C, S*^′^, *E*^′^)*/N* (*S*^′^, *E*^′^). For example one could drop the lighter ball, *S*^′^, *N* (*S*^′^, *E*) times, as *E* = *E*^′^. We would then find that *P* (*C*|*S* = *heavy ball, E*) = *P* (*C*|*S*^′^ = *light ball, E*) and, therefore *ε*(*C*|*S, E*; *S*^′^, *E*) = 0, concluding that the “behaviour” of the heavy and light ball were the same, i.e., their responses to the implicit stimulus of the force of gravity is the same.

If instead of a light or heavy ball we chose as *S* a cat: What response variable should we choose and what is a reasonable null hypothesis? If we choose as state variable its centre of gravity, and response variable the time from falling to impact, then, in an ensemble of *N* (*S, E*) events where the cat fell, we can count the number of events, *N* (*C, S, E*), associated with the response *C*. What should we choose for *S*^′^? If we measure response to falling, *C*, by the changes in the centre of gravity and have an ensemble of observations of the objects, then we will find that *P* (*C*|*heavier ball, E*) = *P* (*C*|*lighter ball, E*) = *P* (*C*|*cat, E*). We can choose any as our subject *S*, and any other as our benchmark *S*^′^, concluding that there is no difference in the responses to the stimulus between any of the systems and therefore their response should not be considered a behaviour. However, if we choose as response — “landing feet first” — this is associated with particular state variables — legs/feet — and so using a ball as a benchmark is nonsensical. In this case a useful benchmark would be some other animal with legs, preferably 4, such as dogs. In this case we would find for *C*, defined as landing feet first, that *P* (*C*|*S* = *cats, E*) ∼ 1 and *P* (*C*|*S*^′^ = *dogs, E*) ∼ 0, thus distinguishing falling cats from falling dogs and allowing us to conclude that landing feet first is a behaviour according to our operational characterisation. So, how can these examples be understood as “choices”? Choice here appears as a consequence of using an ensemble where different responses to a stimulus are observed between a system and a benchmark. Another benchmark is associated with the evolutionary origin of the behaviour, as a response to the stimulus of natural selection when the ancestors of cats gradually passed to an arboreal existence.

#### 2.5.1 A Bayesian Perspective on Behaviour

Having shown how to characterise behaviour in terms of an external ensemble of events we will now turn to how it is characterised in terms of an internal ensemble, as it is precisely this that is the principal source of ambiguity and difficulty in defining behaviour across disciplines. Using the examples of [3], as shown in the Supplementary Material, in spite of the fact that each of the examples refers to a specific event, such as “A spider builds a web”, behind them all is the mental model of an ensemble of events. In other words, no behavioural biologist when considering whether “A spider builds a web” is a behaviour is thinking of a specific spider building a specific web. Rather, the mental model is that of a class of subjects — “spiders” — distinct to other classes of subjects, and a class of state changes in the environments of the subjects — “no webs → webs”. As emphasised, a biologist will not be thinking that all spiders build webs, and even less so that, in general, other animals build webs. Thus, the mental model implicitly compares spiders to other classes of similar subjects that don’t build webs in concluding that spiders building webs is a distinguishing behaviour.

However, we have argued that a necessary condition that a stimulus and a response be a behaviour is that there are multiple potential responses to the same stimulus. If this is the case, then *P* (Δ_*i*_|*R*) *<* 1 for any particular response *i* to the stimulus *R*. However, when we consider spiders building webs or cats falling feet first, we generally consider these to be instinctual behaviours rather than learned behaviours, with *P* (Δ_*i*_|*R*) ∼ 1, wherein the internal ensemble considers all web-building spiders that build webs and all cats that fall feet first. In this sense, the behaviour is universal and there is no “choice” with respect to ensembles that are restricted to web-building spiders or falling cats. However, it is the fact that these behaviours are not truly universal, when considering a wider ensemble that permit us to consider them as behaviours that link particular stimuli to particular responses. Thus, extending the ensemble associated with state changes in the environment corresponding to webs to subjects other than web-building spiders then *P* (Δ_*i*_|*R*) *<* 1. For example, an internal ensemble of none-web-building spiders and web-building spiders allows us to distinguish the behaviour of those spiders that build webs from those that do not. Similarly, an internal ensemble of other animals that also fall in a gravitational field, allows us to distinguish the behaviour of cats as being different to those of other animals.

When we ask *why* spiders build webs, or cats fall feet first, we also have in mind another internal ensemble, one associated with the evolutionary history of the animals. So, to understand the behaviour of spiders building webs, we can envision a time in an evolutionary past where there were no web-building spiders, and then a later time when there were, with a corresponding advantage and corresponding natural selection. In this sense the behaviour is interpreted as an adaptation. This is also true for cats landing feet first, where the implicit comparison could be with evolutionary ancestors of cats that did not possess this reflex but later developed it, or with other animals that did not experience the same evolutionary pressure from natural selection to develop it. Thus, the “choice” element here is implicit in the idea of considering internal ensembles beyond a specific subject and environment.

The implicit idea of an internal ensemble also offers an explanation of why the case from S! of a beetle being swept along by a current was not considered by the vast majority of respondents as a behaviour. Just as in the case of falling balls and cats, where we would not interpret the state changes associated with their centre of gravity in response to the stimulus of gravity as a behaviour, so the physical response of a beetle to a strong current is viewed as a universal response to a physical stimulus analogous to gravity.

The take home message from this section is that, in distinction to an external ensemble, where we can, in principle, begin with a zero prior-knowledge state, when using an internal ensemble the characterisation of a behaviour is subject to a large number of biases that depend on discipline and degree of expertise in the discipline, among other factors.

### 2.6 Characterising Behaviour

Summarising the above discussion, a behaviour characterised in terms of en external ensemble of observations requires the following components:

1. A system, *G* = (*S, E*), divided into a subject, *S*, and an environment, *E*.
2. A set of state variables for the system, **X**(*G, t*), divided into state variables, **X**(*S, t*) and **X**(*E, t*).
3. A (potentially unobserved or unobservable) stimulus, *R*(*t*).
4. A set of potential responses, Δ, measured as observable changes, Δ**X**_*R*_, in a subset, **X**_*R*_ ⊂ **X**(*G, t*).
5. An ensemble of *N* (*S, E*) observable events, 𝒱, and a subset, 𝒱_*C*_, of *N* (*C, S, E*) co-occurrences, *C* = (Δ, *R*), of stimulus and response, or occurrences, *C* = Δ, of a response in the system *G*.
6. Probabilities *P* (*C*|*S, E*) = *N* (*C, S, E*,)*/N* (*S, E*).
7. A benchmark system, *G*^′^ = (*S*^′^, *E*^′^), where *C* can also be observed.
8. Probabilities *P* (*C*|*S*^′^, *E*^′^).

As a given stimulus may not change the state variables that have been chosen to measure a potential response, *N* (*C, S, E*) will depend on the particular state variables chosen.

Using the above conditions we can then characterise a behaviour through the following:

*A behaviour for a subject, S, in an environment, E, is characterised by a statistically significant difference between, P* (*C*|*S, E*) *and a relevant benchmark P* (*C*|*S*^′^, *E*^′^), *where C is a co-occurrence*, (Δ, *R*), *of a stimulus R and a response* Δ, *or an occurrence of* Δ.

We claim that if the above set of requirements are satisfied with an appropriate dataset, then our definition of behaviour is completely operational. Of course, different benchmarks can lead to different results. However, we believe that this is an advantage, rather than a disadvantage, of our approach.

### 2.7 Conduct versus Behaviour

We have presented an operational characterisation of behaviour, and/or behaviour change, based only on a statistical ensemble of observations, that is applicable to any discipline or domain where behaviour is a relevant concept. However, this characterisation does not address the question of: How do we predict behaviour? which is related to the question: Why does a certain behaviour occur? With these questions in mind we make a distinction between “behaviour” and “conduct”, as the etymology of the latter links to the reasons why a certain behaviour is exhibited. For many behaviours the associated “why is very often, implicitly if not explicitly, evolution. Thus, when hunger-hormone ghrelin levels are high (cause-stimulus) we may forage for food (effect-response), with the stimulus-response pair, quantified probabilistically on a suitable ensemble of observations, constituting the behaviour [13]. However, it is the evolutionary pressure from natural selection that has gradually guided individuals to forage for food when hungry that provides the “why — the ultimate, as opposed to proximate, cause — behind this behaviour.

From a human perspective, if we think of behaviours as being the actions a person demonstrates, then conduct can be thought of as the relationship between the action and the normative environment in which the behaviour is exercised. This normativity can be biological in nature eating, reproduction etc. or cultural, where, for example, a particular behaviour might be acceptable in one culture but not in another. Although the complexity of human conduct is enormous, there are certain behaviours that are fundamental and are in common with other organisms [14]. Beyond the biological, as emphasised, human conduct is at the heart of the vast majority of social, health and environmental problems that we face. When such problems are viewed from a behavioural standpoint, we tend to speak of behaviour change [15]. However, without a more thorough understanding of why a certain behaviour is exhibited, it is impossible to predict how easily it may be changed as we have no idea about the normative restrictions that our biological and cultural evolution have imposed. Thus, it is more appropriate to speak of conduct change, as this accounts for the fact that we must distinguish between individual and normative environment and must understand the why behind the behaviour.

Thus, in the rest of the paper we will refer to conduct and conduct change as opposed to behaviour and behaviour change to emphasise the fact that we wish to understand why a certain behaviour is adopted and/or changed. A consequence of this change of perspective is that we must be able to predict conduct and conduct change as it is the determination of those factors that lead to a certain behaviour being adopted that will allow us not only to better understand its origins, but also to determine to what degree it is changeable, i.e., its degree of plasticity

## 3 Constructing a Theory versus a Model for Conduct and Conduct Change

Our characterisation of behaviour up to now has been purely descriptive and has not addressed the question of: How do we predict it? which is related to the question: Why does a certain behaviour occur? We have emphasised that identifying and quantifying a behaviour is a statistical inference problem, and therefore so is its prediction. This statistical inference problem can be addressed using an objective, external ensemble of observations, 𝒱^*e*^, or a subjective, internal ensemble, 𝒱^*i*^. In this latter case, *P* (*C*|*S, E*) represents, in a Bayesian sense, a degree of belief about the relation between *C* and *S* and *E*. Thus, when a behavioural biologist makes the assertion that: A spider building a web is a behaviour — they are building a classification model with a degree of belief that spiders and webs are linked. They are not necessarily saying that a spider is a sufficient condition for a web, i.e., *P* (*web*|*spider*) *<* 1, as there are spiders that don’t build webs. Neither are they saying that a spider is a necessary condition, i.e., *P* (*spider*|*web*) *<* 1, as there are webs that were not built by spiders. The Bayesian classification model *P* (*C*|*S, E*) also serves as a prediction model so that, for example, the next time the biologist sees a web they will predict with a high degree of confidence that a spider will be close by, i.e., co-occur with the web. There is, however, no underlying theory associated with the models, rather, it is a heuristic based on observation and experience.

In the case of an external ensemble, we might think that, as *P* (*C*|*S, E*) = *N* (*C, S, E*)*/N* (*S, E*), we can just count the number of co-occurrences of *C, S* and *E*, and of *S* and *E*. To do this, however, we need to specify both *S* and *E*, using the state variables **X**(*S, t*) and **X**(*E, t*). Taking our falling cat example, what state variables should or could be taken? Cat shape? Cat size? Cat age? Cat sex? Cat breed? Cat genetic mutations? The list is potentially endless. Do we restrict our ensemble to only cats? It is known that some other species exhibit similar behaviour. And what about the environment? Falling from different heights? Falling from different objects? Falling at night versus day? Falling in a strong wind? And if this sounds somewhat pedantic, think about an important health-related conduct such as smoking, or overeating. What do these depend on? Person’s age? Person’s sex? Person’s occupation? Person’s income? Parent’s habits? Person’s personality? Person’s physiology? Person’s genetics? etc. etc. etc. And the environment? Cigarette or food availability? Price? Quality? Peer pressure? Again, the list is truly endless. There are two important points here: the probability for a behaviour is multifactorial, depending on many different subject and environment variables, which, generally, cover many different spatial and/or temporal scales and, therefore, multiple disciplines. In principle, we should consider a probability distribution over the space of all possible subjects and environments. Of course, this is completely infeasible. So what subject and environment variables should be considered? First we should ask what state variables are included in the characterisation of the behaviour itself. In the case of falling cats landing feet first, it makes more sense to restrict to “similar” subjects. This could be similar along the dimension *X*_1_(*S, t*), that could refer to the types of animal considered. So, if *X*_1*j*_ (*S, t*) = *X*_1*j*=*cat*_(*S, t*) we consider only cats as our observed subjects. On the other hand, we could consider a wider range of animals to try to determine what physiological or anatomical features of cats differentiated them from others in their ability to land feet first.

Certainly one can imagine an ensemble of controlled experiments of cats and other animals where the behaviour is observed. Suppose it was found that *P* (*C* = *land feet first*|*X*_1*j*=*cat*_(*S*)) = 0.95 and *P* (*C* = *land feet first*|*X*_1*j cat*_(*S*)) = 0.05. Given a certain ensemble size we can use equation (1) to determine that the two groups behave differently. We might then wonder why *P* (*C* = *land feet first*|*X*_1*j*=*cat*_(*S*)) =*/* 1? Why don’t cats always land feet first? and Why do some other animals land feet first? To answer these questions we must propose other measurable state variables as possible sources of variation. For example, if state variable *X*_2_(*S, t*) refers to a cat’s age, then *X*_2*i*=*age<*8*weeks*_(*S, t*) could refer to those cats that are less than 8 weeks old and *X*_2*i*=*age>*15*years*_(*S, t*) to those cats older than 15 years. We can then compare the behaviour of older and younger cats to determine if the ability to perform the behaviour is age dependent [20]. We could also perform controlled experiments to determine if the environment variable, *X*_1_(*E, t*), referring to the height from which the cat fell made a difference. For example, with *X*_1*j*_ (*S, t*) = *X*_1*j*=*<*1*m*_(*S, t*) to determine the probability of landing feet first when dropped from a height of less than 1m. In a similar fashion we can imagine a host of other controlled experiments, where subject state variables and environment state variables are varied and *P* (*C*|**X**(*S, t*), **X**(*E, t*)) calculated. If we imagined for simplicity that each state variable only took values, (0, 1), then, for *N*_*S*_ subject state variables and *N*_*E*_ environment variables, there are 2^*NS*+*NE*^ different system states that must be experimentally examined in order to determine how a conduct depends on these variables. This is a manifestation of the curse of dimensionality, where the standard response is: “divide and conquer”, i.e., to perform separate experiments on only a small subset of state variables.

Note that *P* (*C*|*X*_*i*_), *P* (*C*|*X*_*j*_) and *P* (*C*|*X*_*i*_*X*_*j*_) represent different prediction models for the potential conduct *C*. We can estimate these probabilities on a given ensemble, 𝒱_*tr*_, the “training ensemble”, and then apply them as predictions on another previously unseen ensemble 𝒱_*te*_, the “test ensemble”, to predict for a given subject in a given environment the probability with which it will exhibit the conduct. We have emphasised that Newton’s Second Law applied in the presence of a uniform gravitational field is a theory that both predicts and explains the trajectory of the centre of gravity of both balls and cats. We have further emphasised that it is manifestly *not* a theory that predicts that falling cats will land feet first. And if it does not predict then it cannot explain. Does the sophisticated treatment of **??**, that models a cat as two cylinders (back and front part of the cat) and connected by a flexible spine, represent a theory? If the conduct depends on any state variables, such as age, size etc. that are not contained in this model then it cannot predict and therefore, again, cannot explain the dependence on these other variables. In reality, this treatment represents a mechanical model that just explains why the cat’s behaviour does not violate any physical laws. Neither Newton’s Laws nor gauge theory of deformable bodies can predict or explain the vast majority of the countless other conducts and observables associated with cats or, indeed, any living system. As a theory, Newton’s Second Law is universal in that it predicts that, given a force *F*, independently of its origen or details, then the object on which the force is applied will accelerate at a rate *a* = *F/m*. Thus, all forces, of a given magnitude and direction, viewed as stimuli, will result in the same response. Newton’s Second Law also “explains” the relation between stimulus, as a proximate cause, and response. However, it does not explain the origin of the force itself; there are many phenomena that can lead to a force of a given magnitude and direction. Moreover, as emphasised in the cat example, it only explains system responses as measured by particular subsets of state variables, such as the centre of gravity. There are many responses, modelled by state changes to stimuli, that are not predicted or explained by Newton’s Laws, or, indeed, any other physical laws. However, these responses must, of course, be consistent with those physical laws.

Darwin’s theory of evolution is another relevant theory that we can appeal to for prediction and explanation in biological systems. Unfortunately, although Darwin’s theory, in principle, applies to both Ω(*S, E*) and *u* (*S, E*), we cannot use it to predict which *U*_*i*_ will be selected, as we do not have an adequate representation of 𝒰 (*S, E*). Indeed, generally, neither do we have an adequate representation of Ω(*S, E*).^3^ What Darwin’s theory can do however, as we will argue, is provide a theoretical framework for explaining predictions found by other means.

## 4 Constructing a Theory for Conduct and Conduct Change using Bayesian Classifiers

As conduct is to be predicted and explained through estimation of *P* (*C*|*S, E*) = *P* (*C*|**X**(*S*)**X**(*E*)) then its prediction is a problem in predicting class membership. Thus, we wish to build prediction models for predicting the probability of occurrence of *C* given a particular subject and/or environment. When the size of the state space, |Ω(*S, E*)|, is small, then *P* (*C*|**X**(*S*)**X**(*E*)) = *N* (*C, S, E*)*/N* (*S, E*) can be estimated directly in frequentist terms without excessive sampling error. However, when |Ω(*S, E*)| is large then *N* (**X**(*S*)**X**(*E*)) = 0, 1. This is just the standard curse of dimensionality, and a statistical or machine learning model must be used. *P* (*C*|*S, E*) can be approximated directly as a discriminative classifier using, for example, logistic regression, neural networks, or decision trees, and their more sophisticated variants, such as random forests or xgboost There are, however, drawbacks to a purely discriminative approach, as without knowledge of the distribution *P* (**X**(*S*)**X**(*E*)), the classifier cannot be used to generate new data points. Additionally, without *P* (**X**(*S*)**X**(*E*)), we do not know how frequent is a particular state of the subject and/or environment. Thus, we may have a strong predictor of a given conduct, but that it is rare among the subjects and/or environments where the conduct is observed.

With this in mind we advocate a generative approach using Bayesian Classifiers [21], constructed via Bayes Rule

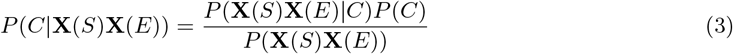

where *P* (*C*) is the prior probability of observing the conduct *C, P* (**X**(*S*)**X**(*E*)|*C*) is the likelihood function that adjusts the estimated probability for *C* given the data **X**(*S*) and **X**(*E*), *P* (**X**(*S*)**X**(*E*)) is the evidence and plays no role in the assignation of a class to an observation in the ensemble but does affect the estimate of the posterior probability *P* (*C*|**X**(*S*)**X**(*E*)). From the frequentist perspective, the natural arena for equation (3) is an ensemble, 𝒱, of *N* observations, of which in a sub-ensemble, 𝒱(*S, E*), there are *N* (**X**(*S*)**X**(*E*)) observations where there is a co-occurrence of the subject state variables **X**(*S*) and the environment state variables **X**(*E*). Similarly, there are *N* (*C***X**(*S*)**X**(*E*)) co-occurrences of *C*, **X**(*S*) and **X**(*E*) in the same sub-ensemble. An important feature of Bayes rule is that it can be naturally iterated as we add or remove information. For example

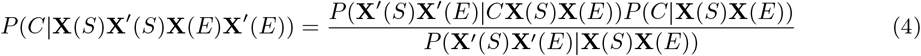

represents what happens when, from an ensemble of *N* events, *P* (*C*|**X**(*S*)**X**(*E*)) is calculated based on the state variables **X**(*S*) and **X**(*E*), then new information is added in the form of observations of new state variables **X**^′^(*S*) and/or **X**^′^(*E*). In this setting, a Bayesian model of the form *P* (*C*|**X**(*S*)**X**(*E*)) is first proposed by making observations of the co-occurrences of *C*, **X**(*S*) and **X**(*E*) in the ensemble 𝒱_*tr*_ of *N* events, which forms our training set. This model can be then tested and its performance measured on a test ensemble, 𝒱_*te*_. We can then ask if the addition of new information, by including state variables **X**^′^(*S*) and **X**^′^(*E*), distinct to those included in **X**(*S*) and **X**(*E*), can lead to a better model. Note that in this case we are using the same ensemble 𝒱 of *N* observations, but incorporating new information from that ensemble.

Taking the example of section 3 for falling cats, we could begin with a model *P* (*C*|*X*_*i*_) - Theory 1, where *C* is landing feet first and *X*_*i*_ is the cat’s age. We can then add in information *X*_*j*_ associated with the height of the fall and consider the model *P* (*C*|*X*_*i*_*X*_*j*_) - Theory 2, which is related to Theory 1 via Bayes rule

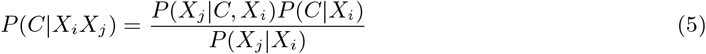

The question is now: Is Theory 2 better than Theory 1? We can consider two principle requirements: Does it predict better? Does it explain better? The former can be addressed objectively through one or more, well known performance criteria. It is independent of the semantics of the state variables chosen, i.e., we can predict without understanding why a prediction is good or bad. The latter is more subjective and susceptible to disciplinary bias. In the present example, if we find that very young cats are found to be less probable to exhibit the conduct then we need to explain why. For example, we might observe that the probability increases with age between 2 weeks and 10 and use that to understand that it is a reflex that is developed. We might also observe that older cats are less able to land feet first and wonder why. Perhaps because older cats tend to be heavier — this can be tested by adding a third state variable *X*_*k*_ that is weight — or it could be because older cats are less flexible. This also can be tested with other state variables. If we also found that the probability decreases substantially when the fall height is below a certain threshold, then we might explain that by determining that the cat needs a certain amount of time in order to contort its body in order to land feet first. Thus, we may conclude that, both in terms of predictability and explainability, Theory 2 is a better theory of falling cats than Theory 1. Of course, there are many other “theories” that could be proposed by including in other potential factors that influence the probability of observing the conduct. Each can be tested for its predictability and explainability. Of course, it can occur that adding in more state variables does not lead to a better theory. Indeed, it may be better to remove some. In principle, this can be done by inverting equation (5). However, it is more insightful to think of Theory 2 in terms of the joint probability *P* (*CX*_*i*_*X*_*j*_). Passing to Theory 1 can be done by considering 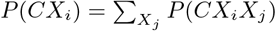, i.e., marginalising on *X*_*j*_ .

The construction of a theory based on the observations associated with an external ensemble is completely dependent on the measurements made. For example, if the original ensemble contained measurements of *X*_*i*_, but not *X*_*j*_, then it would not be possible to marginalise on *X*_*j*_ and then compare the two theories, or indeed any other containing *X*_*j*_, using equation (5). What theories are available to construct and compare from a given ensemble therefore depends on the choice of state variables measured. In other words, we can imagine Bayesian priors 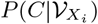 and 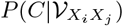, where by 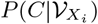 we mean the probability for observing the conduct on an ensemble of events where in those events only *X*_*i*_ was measured, while by 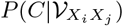 we mean the probability for observing the conduct on an ensemble of events where in those events both *X*_*i*_ and *X*_*j*_ were measured in each event. In this sense, when we write *P* (*C*|*X*_*i*_) we could mean 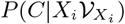, as calculated on an ensemble where only *X*_*i*_ was measured, or 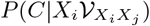, the conditional probability for *C* given a value for *X*_*i*_ as obtained from an ensemble where both *X*_*i*_ and *X*_*j*_ were measured. From the latter we can compute *P* (*C*|*X*_*j*_) and *P* (*C*|*X*_*i*_*X*_*j*_) but from the former we cannot. In fact, this example illustrates perfectly the effect of disciplinary bias, manifest as a Bayesian prior, not on the choice of state variables that enter into the prior but, rather, on the choice of ensemble in the first place. Thus, for example, if *X*_*j*_ is an important variable then, although a choice of prior, 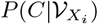, might be improved upon by calculating the likelihood 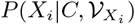 to find, using Bayes rule, a better model, 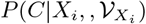, this choice of prior ensemble does not permit the inclusion of *X*_*j*_ . However, a choice of prior ensemble, 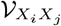, allows various routes to finding a better model, first using a prior 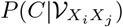, and incorporating joint information on *X*_*i*_ and *X*_*j*_ via the likelihood 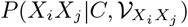 to find, again using Bayes rule, the model 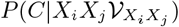, or secondly, by choosing as prior, 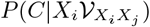 or 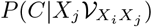 then calculating the likelihoods 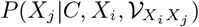 and 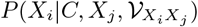 respectively to find 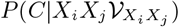.

Again, lest this be thought as being somewhat pedantic, it is exactly this that is one of the principal reasons why the major problems associated with human conduct prove so intractable. For example, in the case of obesity, where the conducts of excess consumption and sedentariness are key causes, a multifactorial approach to the problem is impeded by the disciplinary biases that lead to data being collected in a given study for only a narrow subset of state variables; for instance psychological variables in one study on one population, versus socio-economic factors on another. As emphasised, such studies are incommensurable, in that they cannot be combined to analyse the potential interactions between the different factors. However, if psychological variables and socio-economic variables are collected from the same population, then they can, in principle, be combined. We would argue that the design and execution of experiments on a particular population that only include a narrow subset of state variables associated with a particular discipline is a consequence not only of our biased mental models, but also of an incapacity to fully comprehend and then operationalise the high degree of multifactoriality inherent in complex adaptive systems.

## 5 Approximating the Bayesian Classifier

The Bayes classifier is the classifier having the smallest probability of misclassification of all classifiers that use the same set of features. However, this conclusion is based on being able to calculate *P* (**X**(*S*)**X**(*E*)|*C*). Unfortunately, due to the curse of dimensionality, an exact calculation is not feasible. Although there are different types of approximation, here we will advocate one that assumes that *P* (**X**(*S*)**X**(*E*)|*C*) can be factorized

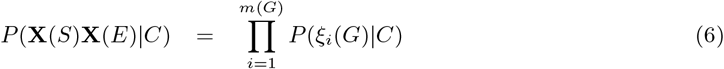

where **X**(*G*) is partitioned into a set of *m*(*G*) non-overlapping blocks [22]. For instance, for two subject variables, *X*_1_(*S*) and *X*_2_(*S*), and two environment variables, *X*_1_(*E*) and *X*_2_(*E*), some potential partitions are: *{X*_1_(*S*)*}{X*_2_(*S*)*}{X*_1_(*E*)*}{X*_2_(*E*)*}, {X*_1_(*S*), *X*_2_(*S*)*}{X*_1_(*E*), *X*_2_(*E*)*}, {X*_1_(*S*), *X*_2_(*S*), *X*_1_(*E*)*}{X*_2_(*E*)*}* and *{X*_1_(*S*), *X*_2_(*S*), *X*_1_(*E*), *X*_2_(*E*)*}*. The first partition corresponds to the well-known Naive Bayes approximation (NBA) [23], where the four state variables are assumed to be independent when conditioned on the class *C*. By considering variable combinations we are accounting for potential interactions between them. However, even though there are strong interactions, this does not necessarily imply that the NBA is bad, as there are always cancelations in the errors between different variables and variable values [22].

In order to calculate *P* (*C*|**X**(*S*)**X**(*E*)), we use Bayes rule and, to avoid an explicit calculation of the evidence function, we consider

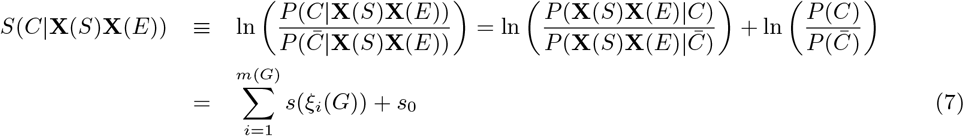

where 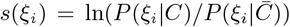 is the weight of evidence [24] associated with the combination *ξ*_*i*_ and 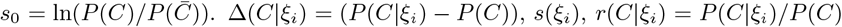 and *r*(*ξ*_*i*_|*C*) = *P* (*ξ*_*i*_|*C*)*/P* (*ξ*_*i*_) can all be thought of as measures of the effect size of the variable combination *ξ*_*i*_ for observing the conduct *C*, with *r*(*C*|*ξ*_*i*_) and *r*(*ξ*_*i*_|*C*) representing the degree of sufficiency and necessity of the combination *ξ*_*i*_ for observing *C*. An important feature of this methodology is that the calculation of all relevant quantities is based only on event counts: *N*, *N* (*ξ*_*i*_), *N* (*C*) and *N* (*Cξ*_*i*_). For example, *P* (*ξ*_*i*_|*C*) = *N* (*Cξ*_*i*_)*/N* (*C*). Thus, *S*(*C*|**X**(*S*)**X**(*E*)), or any other relevant function, can be built from only *N*, *N* (*ξ*_*i*_), To convert these counts into reliable probabilities we require that all state variables are categorical with a finite number of categories. In this case *N* (*ξ*_*ij*_) represents the number of events in the category *j* of the variable *ξ*_*i*_. More categories means higher resolution to analyse the relation between *ξ*_*i*_ and *C*, but also means a smaller statistical sample in a given category, thus allowing for sampling errors leading to misclassifications. The coarse graining associated with the division into categories is an important element in the construction of a prediction model that can affect its performance.

If *S*(*C*|**X**(*S*)**X**(*E*)) *>* 0, then subjects described by the state variables **X**(*S*), in environments described by the state variables **X**(*E*), are predicted to exhibit the conduct, and vice versa when *S*(*C*|**X**(*S*)**X**(*E*)) *<* 0. Besides pure classification however, *S*(*C*|**X**(*S*)**X**(*E*)) can be used to reconstruct *P* (*C*|**X**(*S*)**X**(*E*)) as a probability distribution by inverting equation (7). to find

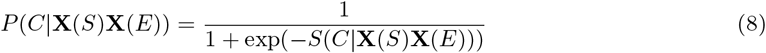

The predictability of the full model can be gauged in standard fashion by deriving it first on a training ensemble then applying it to a test ensemble and measuring the model’s performance. If we consider pure classification, this can be done by using a standard measure, such as a confusion matrix or a ROC curve. If we are considering *P* (*C*|**X**(*S*)**X**(*E*)) then we can use, for example, *δ* = _*i*_ |*P* (*C*|*S*_*i*_*E*_*i*_) − *P*_*o*_(*C*|*S*_*i*_*E*_*i*_)|, where the sum is over the events in the ensemble and *P*_*o*_(*C*|*S*_*i*_, *E*_*i*_) is the observed probability. In the case where the subject and environment specification is sufficiently detailed then *P*_*o*_(*C*|*S*_*i*_, *E*_*i*_) = 0, 1, as every event is unique.

In terms of predictability there are three steps that can strongly influence model performance: i) the choice of measurements to be made in 𝒱_**G**_; ii) the choice of which variables *X*_*i*_ to combine in *ξ*_*i*_; iii) the choice of which *ξ*_*i*_ to include in the final model. Step i), seen as a Bayesian prior, is the most important, and reflects the negative impact of disciplinary bias. For example, to answer the question: Is factor A more important than factor B in predicting a conduct, or indeed any dependent variable, an ensemble must be constructed where both A and B are collected. Although this may seem an obvious point, the lack of data sets that are multiscale and multidisciplinary is evidence that this point is not incorporated as a focal element in scientific planning. Even when there exist important exceptions to this, such as the UKBiobank [25] or the All of Us program [26], this does not mean that models built with this data are multiscale and multi-disciplinary. Step ii) can be addressed by analysing interaction measures, such as Δ(*X*_*i*_*X*_*j*_) = (*P* (*X*_*i*_*X*_*j*_)|*C*) − *P* (*X*_*i*_|*C*)*P* (*X*_*j*_ |*C*)), where, if Δ(*X*_*i*_*X*_*j*_) is statistically significant, then a new variable *ξ*_*k*_ = *X*_*i*_*X*_*j*_ should be introduced. Step iii) is the standard problem of feature selection. There are different methods for selecting features to be included in a model. Here we will use a filter method, using the effect size of *ξ* as a means to rank the features. For instance, by keeping those features that have the highest score *s*(*ξ*). However, *s*(*ξ*), is based on probabilities, and is therefore independent of ensemble size. This means that a feature might be erroneously included due to sampling error. To avoid this we can use a statistical diagnostic, such as the binomial test of equation (2), to evaluate the statistical significance of the effect size. For example,

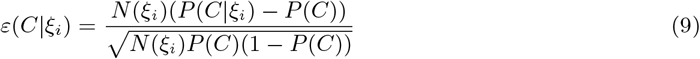

where Δ(*C*|*ξ*_*i*_) = (*P* (*C*|*ξ*_*i*_) − *P* (*C*)), is statistically significant at the 95% confidence interval when |*ε*| *>* 1.96 and the binomial distribution can be approximated by a normal distribution. Additionally, it is important to be able to measure and compare the coverage of a factor *ξ*_*I*_, as measured by *N* (*ξ*) or *P* (*ξ*_*I*_). Equation (9) represents a single quantity that captures both the effect size of *ξ*_*i*_, through its dependence on Δ(*C*|*ξ*_*i*_), as well as its coverage through *N* (*ξ*). The combination of effect size, coverage and statistical reliability give a quite complete representation of the contribution of a given combination of subject and environment variables to the prediction of the conduct or conduct change.

The fact that predictions can be broken down into predictive “blocks”, *ξ*, that contain relatively few state variables (just one in the case of the NBA), means that the models are transparent and therefore “explainable”, in that the contribution of each variable combination is manifest in terms of its effect size, its coverage and its statistical significance. These factors also go a long way to providing an explanation for the predictions by giving clear and transparent quantitative and objective indicators that permit the evaluation of the role of each factor. Unlike Shapley scores [27], which have been extensively promoted, and also criticised for being potentially misleading [28], feature selection using *ε*, or effect size directly, is a filter method, and therefore independent of the ML algorithm used. This feature selection method is particularly adapted to the factorised Bayesian classifier algorithm we are advocating. It is also an absolute not relative measure of feature importance, as it directly incorporates the statistical significance of the feature and changes as a function of sample size.

Although explainability is facilitated by being able to examine each predictive feature combination separately, it also requires an understanding of the meaning of *ξ*_*i*_ as a predictor of *C*. This understanding is a property of Human, as opposed to Artificial, Intelligence, and principally involves trying to fit the observed statistical relations into our own mental model of the phenomenon. This is a delicate procedure, especially in the context where the conduct is dependent on a host of predictors that are linked to very different disciplines. For example, it might be difficult for a psychologist to intuit why a particular mutation in a particular gene could influence a conduct, whilst a geneticist might be puzzled trying to understand the impact of self-efficacy on a person’s decision making. This is not problematic when the ensemble studied by the psychologist only contains psychological state variables, and that studied by the geneticist only genetic data. However, the search for the best model should contain both variable types. One could rely on statistical analysis used in the feature selection procedure only, but then one could end up “p-fishing”. It is essential to develop an understanding of the possible causal nature of any relationship between *ξ*_*i*_ and *C* and, potentially, between one *ξ*_*i*_ and another. Is *ξ*_*i*_ a direct cause or an indirect cause? Are there confounding factors? Furthermore, if we are speaking about conduct change, it is vital to know which features are susceptible to an intervention, i.e., are they actionable? The optimal combination between Artificial Intelligence and Human Intelligence necessary to produce a model that is both predictive and explainable we can term Hybrid Intelligence [29].

### 5.1 A health-related application

As examples of our framework, as it relates to the characterisation of behaviour, in S1 we present an analysis of the cases considered in [3], analysing them in the context of the properties we have claimed are essential components of a behaviour. In this section, however, we wish to illustrate the discussion and theoretical framework we have presented for predicting conduct and conduct change in the context of a multifactorial real-world example. The data set we will use comes from Project42, associated with the CHILAM laboratory at the Centre for Complexity Science at the Universidad Nacional Autónoma de México (UNAM) and is downloadable there (https://chilam.c3.unam.mx/en/proy-42-eng/data-project42)^4^. A primary objective of the project is to collect multiscale, multidisciplinary data using a diverse set of experimental protocols in order to develop an integrated perspective of the participants with an emphasis on developing predictive and explainable models for obesity and metabolic disease. As two key drivers of these diseases are conductual: nutrition and physical activity, we will use our Bayesian classifier framework to build predictive models for one of these conducts. The specific dataset we consider is associated with an ensemble of 1075 workers, academics and students of the UNAM as subjects. In 2014 multiple protocols were applied to obtain epidemiological, anthropometric, physiological, socio-demographic, socio-economic, social, nutritional, lifestyle and genetic data, comprising more than 3000 data categories (model features), among which were many state variables that represent health-related conducts. The data dictionary and questionnaires used can be downloaded at https://chilam.c3.unam.mx/en/proy-42-eng/data-project42. The Project42 modelling platform (project42.c3.unam.mx) allows for the creation of Bayesian classifiers based on the simplified approximation of the NBA. It permits the user to choose as class, *C*, any of the more than 3000 model features, and as predictors any of the remaining variables. In this particular data set the state variables are principally subject state variables, though there are a number of environment state variables. The data set is associated with an ontology that partitions the variables into categories: Personal data, Personal Background, Family History, Health Self-Assessment, Lifestyle, Health Information, Anthropometry, Nutrition and Laboratory results. The system permits the creation of more than 2^3000^ different models, depending on which variable is chosen as target class and which subset of state variables are chosen as predictors. This allows for the rapid creation and validation of hypotheses with respect to the predictive and explanatory power of a set of chosen predictors. In Table 1 we see the results of a model where the chosen target class was *C* = *no exercise*, representing those participants who reported not participating in any physical exercise at the time of the questionnaire, and the predictors were a set of 58 variables from different categories corresponding to 396 features. The results shown correspond to those features positively correlated with *C* with statistical significance *p <* 0.05 (*ε >* 1.96).

**Table 1:** Showing the 20 potential examples of behaviour as considered in [3].

| Case | Example | Objections | % Approval |
| --- | --- | --- | --- |
| (1) | Ants that are physiologically capable of laying eggs do not do so because they are not queens | (C, G) | 64% |
| (2) | A sponge pumps water to gather food | (B, M) | 74% |
| (3) | A spider builds a web |  | 97% |
| (4) | A rabbit grows thicker fur in the winter | (A, G, I, M) | 13% |
| (5) | A plant's stomata (respiration pores) close to conserve water | (I, K, M) | 38% |
| (6) | A plant bends its leaves towards a light source | (K, M) | 48% |
| (7) | A person's heart beats harder after a nightmare | (B, I, L) | 34% |
| (8) | A person sweats in response to hot air | (G, I, L, M) | 24% |
| (9) | A beetle is swept away by a strong current | (F, M) | 5% |
| (10) | A rat has a dislike for salty food | (B, C, G, J, M) | 66% |
| (11) | A person decides not to do anything tomorrow if it rains | (B, C, G, J, M) | 80% |
| (12) | A horse becomes arthritic with age | (A, B, E, G, M) | 6% |
| (13) | A mouse floats in zero gravity in outer space | (E, F, G, M) | 11% |
| (14) | A group of unicellular algae swim towards water with a higher concentration of nutrients | (F, H, K, M) | 78% |
| (15) | A frog orbits the Sun along with the rest of the Earth | (F, M) | 9% |
| (16) | Flocks of geese fly in V formations | (H) | 99% |
| (17) | A dog salivates in anticipation of feeding time | (B, G, I, M) | 75% |
| (18) | Herd of zebras break up during the breeding season and reform afterwards | (H) | 96% |
| (19) | A chameleon changes color in response to sunlight | (G, M) | 57% |
| (20) | A cat produces insulin because of excess sugar in her blood | (B, G, I, M) | 14% |

**Figure 1:**
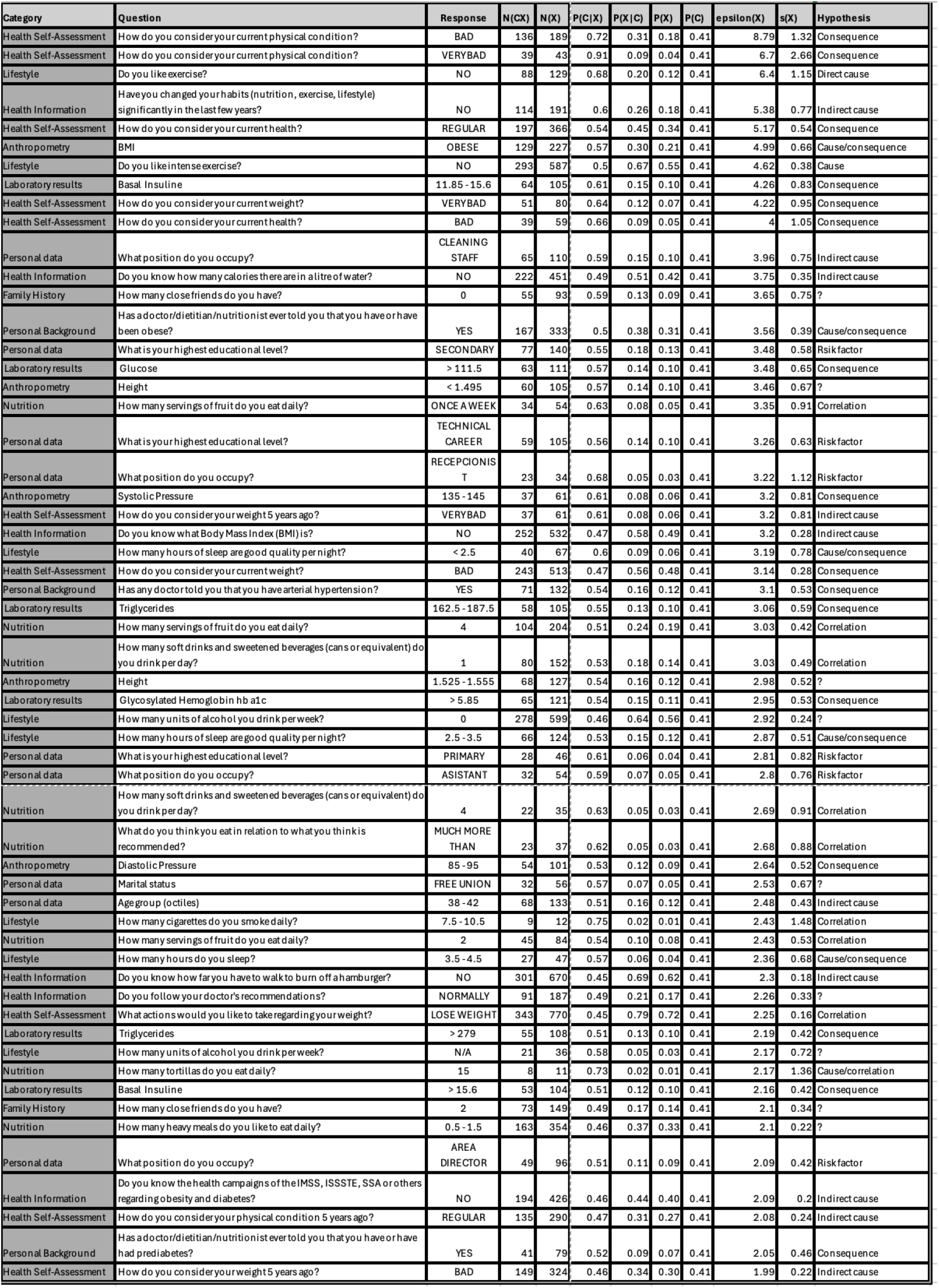
Statistically significant state variables with positive relation with the conduct “no exercise”.

**Figure 2:**
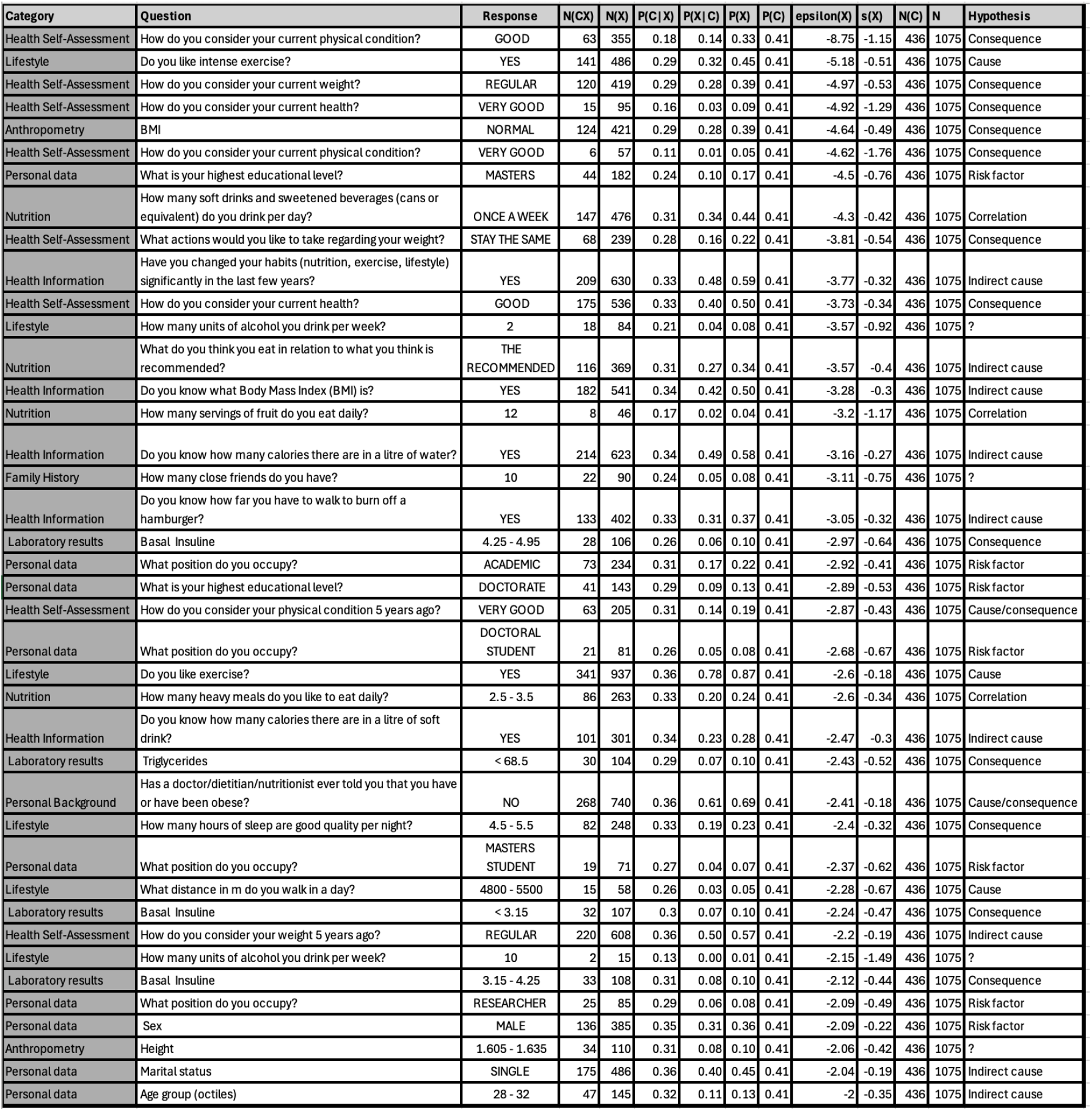
Statistically significant state variables with negative relation with the conduct “no exercise”.

In this example there is no explicit observed stimulus and the response, corresponding to a state change, or rather no state change, is also implicit, based on a self-report of each participant. The null hypothesis in this example is the probability, *P* (*C*), across all subjects for the no-exercise class. *P* (*C*) = 0.41, as *N* (*C*) = 436 of the *N* = 1075 participants reported not doing any physical exercise. To draw an analogy with our previous discussion and examples, consider an ensemble of *N* = 1075 animals falling in a gravitational field. We observe that *N* (*C*) = 436 land feet first. Imagine that we had no prior knowledge about which animals land feet first, but we had data, or could collect data associated with characteristics of the animals to try and predict and understand why certain animals land feet first. If one of the variables was the Linnaean species name then we might observe that 439 of the animals were *Felis catus* and that of them 436 landed feet first. We could then imagine building a prediction model *P* (*C*|*S*), where *S* was a binary variable, *S* = *Felis catus* = *yes, no*. This prediction model would have very good predictive performance and could be used to make future predictions. However, the model would tell us nothing about why or how *Felis catus* was so predictive. Involved in the project we might have some physiologists who could hypothesise that the ability must be related to some physiological adaptations of the animal, such as a highly evolved vestibular apparatus in the inner ear, a very flexible backbone and a lack of a collar bone. Data could be collected to see if *Felis catus* possessed these traits. If this is the case, this provides explainability as to how *Felis catus* lands feet first. We could also see if other animals in our ensemble had any or all of these properties. In fact, we can create prediction models based on any or all of these properties. Although we might make progress on the “how” *Felis catus* lands feet first, by possessing certain adaptations that other animals did not, it would not explain how *Felis catus* obtained these adaptations. If there are some evolutionary biologists in the project, in light of the natural history of *Felis catus*, that evolved from arboreal-dwelling ancestors, the ultimate explanation of why *Felis catus* lands feet first could be proffered, i.e., that natural selection led to a selective advantage for those individuals that developed adaptations that could permit them to land feet first. Thus, we see that the development of a prediction model for this conduct and its interpretation requires not only data, but also a multidisciplinary team to explain the results of the model.

Returning to our present example, we wish to determine those subject and environment factors that best predict and explain why a member of the ensemble is in the no-exercise class. As there is no explicit stimulus measured, there is no natural “no stimulus” benchmark. Hence, we compare the conduct of one group relative to a benchmark associated with another. Here, the benchmark group is associated with an ensemble that is the full set of participants, where the putative behaviour is present with a probability *P* (*C*). Table 1 then shows the results for *P* (*C*|*S, E*), where *S* = *X*_*i*_, as the probability of observing the behaviour in a sub-ensemble of size *N* (*X*_*i*_) defined by the subject variable *X*_*i*_. When *ε*(*C*|*X*_*i*_) *>* 1.96 we can conclude that the conduct of the subjects defined by *X*_*i*_ is significantly different to that of the whole group.

We will comment on the results only to the extent that it illustrates the framework we have presented for predicting and explaining conduct. The first point to make is that of the 396 features 97 (24.5%) are significant at the level *p <* 0.05. Of course, a large number of binomial tests could lead to statistical significance of some features by chance. In this case a correction, such as the Bonferroni correction [30], may be applied. Here, we prefer to show all results to allow for a potential causal and/or actionability interpretation of the observed correlation. Additionally, many of the predictors are significant at far smaller values than *p <* 0.05, e.g., *ε* = 4 corresponds to *p* = 0.00003.

All of the 9 different variable categories contribute predictive features at this significance level, again showing the highly multifactorial nature of the factors that influence the conduct, which, in its turn, demands a completely multi-disciplinary approach with respect to the interpretation of the results. Some features, such as eating a large number of tortillas daily, can have a large effect size (*s* = 2.09), but low coverage (*N* (*X*) = 11), while others, such as if the participant knows what Body Mass Index is, have large coverage (*N* (*X*) = 532), but small effect size (*s* = 0.28). Some factors are more readily identifiable as consequences, such as being obese, or feeling that your current physical condition is very bad, while others, such as not liking exercise, are clearly direct causes, or risk factors, such as low educational level. With the latter, one might wonder why low educational level is a risk factor for this conduct? To answer this, in the online system, one can quickly create a model using a specific educational level as target class to determine what the predictors of this level are which could explain in more causally intuitive terms why low educational level is correlated with sedentariness. In table 2 we show analogous results, but now for the those factors that are correlated with low probability to not exercise, and which are statistically significant at the *p <* 0.05 level, with the results for ordinal variables being the opposite of those that appear in Table 1. For example, we see higher educational level, enjoying intense exercise, good metabolic health and healthy nutrition as being significant factors.

**Table 2:** Showing the 13 potential criteria by which the examples in Table 1 could be judged to be a behaviour.

| Case | Proposed feature | % Approval |
| --- | --- | --- |
| (A) | A developmental change is usually not a behavior | 69% |
| (B) | Behavior is always in response to the external environment | 16% |
| (C) | A behavior is always an action, rather than a lack of action | 14% |
| (D) | All behaviors are directly observable, recordable and measurable | 46% |
| (E) | People can all tell what is and isn't behavior, just by looking at it | 14% |
| (F) | Behavior is always influenced by the internal processes of the individual | 78% |
| (G) | Behavior always involves movement | 8% |
| (H) | Behaviors are always the actions of individuals, not groups | 31% |
| (I) | Behavior is something whole individuals do, not organs or parts that make up an individual | 70% |
| (J) | A behavior is always in response to a stimulus or set of stimuli, but the stimulus can be either internal or external | 87% |
| (K) | Behavior is something only animals (including people) do, but not other organisms | 36% |
| (L) | In humans, anything that is not under conscious control is not behavior | 10% |
| (M) | Behavior is always executed through muscular activity | 18% |

Finally, we can determine the overall predictive power of the model as measured by its ROC curve and the corresponding area, as seen in Figure 3, showing that the model has good predictive power relative to the predictors chosen, as well as being transparent from the point of view of explainability. However, this is just one model chosen from a very large number of possibilities. The advantage of the proposed framework, and the platform itself, is to permit the rapid construction and validation of hypotheses iteratively, using both AI and Human Intelligence combined, to guide the search for predictability and explainability, along with the interpretation and utilisation of the models in real-world situations.

**Figure 3:**
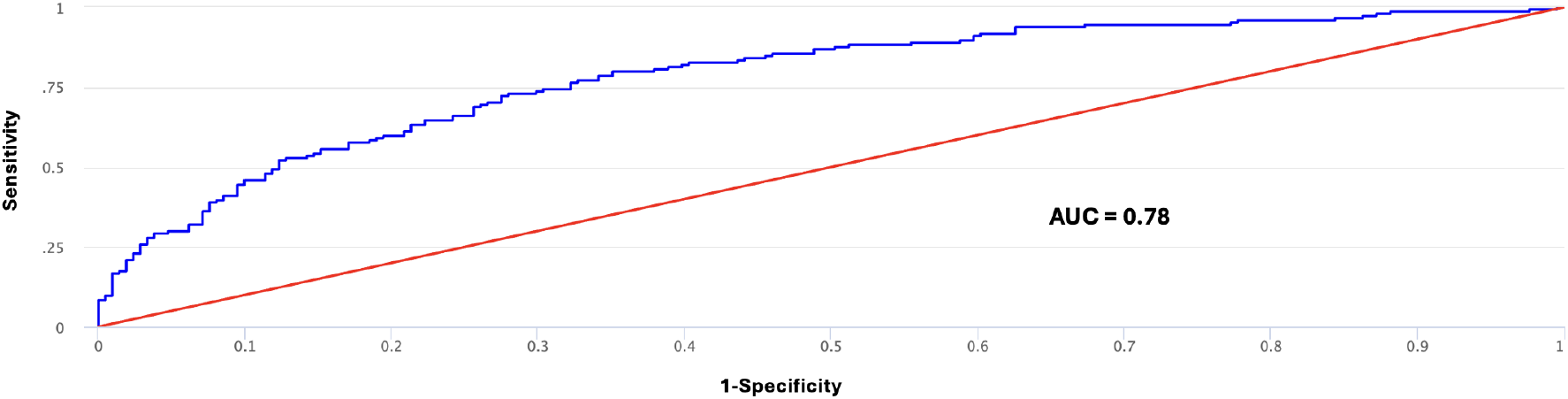
ROC curve for predictive model for *C* = “no exercise” and 58 subject variables.

## 6 Conclusions

In this paper we have presented a general framework for both characterising behaviour as well as predicting and explaining it. We have argued that the wide spectrum of definitions of behaviour, both between disciplines and within them, is a result of the fact that behaviour is seen through a disciplinary lens that acts as a Bayesian prior. To avoid this we presented a characterisation that is operationalised in terms of measurements of stimuli and responses taken from an ensemble of events associated with the subjects of the potential behaviour and their environment. We showed that if our mental models of the behaviour exhibited in a given situation are set aside, then behaviour must be characterised by statistical inference on this ensemble, with *P* (*C*|*S, E*) being the probability for a behaviour *C* for a subject *S* in an environment *E*. We also showed that it is a relative not absolute concept, whereby to distinguish a behaviour in a group of subjects we must compare it to a benchmark group. We also argued that a crucial characteristic of behaviour is that it is associated with multiple potential responses of a system to the same stimulus, or multiple stimuli leading to the same response (multi-causality), and hence was a property of biological systems only. In the Supplementary material we used this characterisation to analyse 20 examples of behaviour in biology that have been used [3] to arrive at a consensus definition, showing how any ambiguity as to what should be considered a behaviour could be resolved.

We then developed a computational framework for predicting and explaining conduct, viewing it as a classification problem, where *P* (*C*|**X**(*G*)) is calculated as a Bayesian classifier using well known machine learning techniques. The set of factors **X**(*G*) represent an approximation to the complete set, **X**_*C*_ — the Conductome — that best predicts and explains the conduct in the context of a particular ensemble. We applied the framework to a highly multifactorial, cross-disciplinary dataset with over 3000 features, deriving an approximation to the Conductome for the conduct “no exercise”, which is well known as a risk factor for obesity and metabolic disease, as a function of 58 state variables containing 396 features. We showed that 25% of these features were predictive at the *p <* 0.05 level, thus showing that this behaviour was significantly correlated with a wide set of predictors.

## Acknowledgments

We are grateful for fruitful discussions with all the members of the Conductome project at the Centro de Ciencias de la Complejidad. This work was partially funded by a SECHITI grant to the Laboratorio Nacional de Ciencias de la Complejidad. CRS is grateful to PASPA UNAM for a sabbatical fellowship and to Antonio del Rio for very fruitful conversations.

## 7 Supplementary Material

In this Supplementary material we will apply our criteria for characterising behaviour to the 20 examples considered by Levitis et al [3] in their analysis of the opinions among, and criteria used by, a group of 181 biologists to characterise behaviour. Table 1 shows the examples used and table 2 the postulated features of behaviours. Thus, for instance, in example (16) of table 1 the potential objection to geese flying in 𝒱 formations as a behaviour is that it violates postulated feature (H); i.e., if we think that behaviour is only a characteristic of individuals, not groups of individuals, then (20) is not a behaviour. Similarly, example (4) violates 4 of the proposed features. The percentage approval figures in both tables represent the percentage of respondents who agreed that the example should be considered a behaviour, or that the postulated feature was an important one for defining behaviour. From table 1 we see that there is an impressive amount of disagreement between behavioural biologists as to what is a behaviour. If we took as a simple requirement that there was agreement at least within 75% of the group as to whether a given example constituted a behaviour or not, then only 12 of the 20 examples reach this level of consensus. If we require 90% consensus, then this number reduces to 6. Further, if we use as a benchmark flipping a coin, then two of the examples, (6) and (19), are indistinguishable from flipping a coin at the *p <* 0.05 level!

In table 3 we see the 20 examples considered by Levitis et al, but in terms of the characterisation of behaviour (conduct) that we have proposed in this paper. The key elements are that there are subjects, an environment, a stimulus, a corresponding response and a benchmark. The key component of our characterisation that cannot be applied here is the concept of an external statistical ensemble of observations. In that sense, *P* (Δ|*R*) or *P* (Δ) should be calculated from an ensemble of observations for each example. Obviously, the examples were chosen based on scientific and experiential evidence associated with the respondent’s mental models of the phenomena. However, there is no explicit associated data sets by which *P* (Δ|*R*) or *P* (Δ) can be calculated. It is notable that in almost none of the examples is there an explicit reference to an environment in which the potential behaviour is taking place. Neither is the presence of an explicit stimulus universal, as is evident in examples (1), (3), (5) and (16). However, the presence of a subject and a response is universal.

**Table 3:** Key elements in the characterisation of the examples of [3] using the definition of behaviour from this paper. NS signifies Not Specified. E+ signifies the hypothesis that the adaptation is selected for, while En - signifies the hypothesis that the adaptation is neutral and E-that it is evolutionarily selected against.

| Case | Subject | Environment | Stimulus | Response | Benchmarks | Adaptation | Behaviour |
| --- | --- | --- | --- | --- | --- | --- | --- |
| (1) | Non-queen ants | NS | NS | Not laying eggs | Queen ants | E+ - division of labour | Yes |
| (2) | Sponge | NS | Food | Pumps water | No food | E+ - food, respiration and waste removal | No |
| (3) | Spider | Web | NS | Build web | Non-web building spiders | E+ - prey capture | Yes |
| (4) | Rabbit | NS | Winter | Thicker fur | Tropical or sub-tropical species | E+ - reduce heat loss | Yes |
| (5) | Plant stomata | NS | NS | Close | Day versus night | E+ - conserve water | No |
| (6) | Plant leaves | NS | Light | Leaves bend towards light | Photonasty | E+ - capture more light | Yes |
| (7) | Person's heart | NS | Nightmare | Beats harder | Non-REM sleep | E+ - processing stressful experiences | Yes |
| (8) | Person | NS | Hot air | Sweats | Primate species that do not sweat | E+ - maintain core temperature | Yes |
| (9) | Beetle | NS | Strong current | Swept away | Other species of beetle | No | No |
| (10) | Rat | NS | Salty food | Dislike | Salt depleted rats | En | Yes |
| (11) | Person | NS | Prediction of rain tomorrow | Decides not to do anything | Persons who go out in the rain anyway | L - | Yes |
| (12) | Horse | NS | Aging | Arthritis | Vertebrates that show little degeneration | E- | Yes |
| (13) | Mouse | Outer space | Zero gravity | Floats | Any other object | No | No |
| (14) | Unicellular algae | NS | Nutrients | Swim towards nutrients | Absence of nutrients | E+ - food acquisition | Yes |
| (15) | Frog | Earth | Gravity | Orbits the Sun | Any other object | No | No |
| (16) | Flocks of geese | NS | NS | Fly in V formations | Non-migratory birds or birds that migrate alone | E+ - energy conservation | Yes |
| (17) | Dog | NS | Feeding time | Salivation | Change feeding time | E+ - production of enzymes for digestion | Yes |
| (18) | Herds of zebras | NS | Breeding season | Break up | Zebra species that don't break up | E+ - mate seeking | Yes |
| (19) | Chameleon | NS | Sunlight | Change color | No sunlight | E+ - camouflage and intra-species signalling | Yes |
| (20) | Cat | NS | Excess sugar in blood | Insulin production | Invertebrates | E+ - glucose storage and usage | Yes |

We have emphasised that behaviour is a relative not an absolute concept, hence we add a benchmark with respect to which *P* (Δ|*R*) or *P* (Δ) can be compared. The benchmark is not unique, as different benchmarks can lead to different conclusions as to whether we would consider the response as a behaviour. This is not a weakness but, rather, an indication that behaviour is relative not absolute, and a motivation to question the mental models by which we consider an example to be a behaviour or not. For some examples, we have chosen benchmarks, i.e., ensembles, where we would not consider the response to be a behaviour, as there are no multiple responses to the same stimulus. However, all the examples, other than (9), (11), (13) and (15), can be thought of as innate responses, linked to an adaptation in the evolutionary past. A choice of benchmark ensemble that considers both ancestors and descendants of the considered subjects, before and after the behaviour was favoured by natural selection, would then lead to a consideration of the responses as being behaviours according to our characterisation. On the contrary, examples (9), (13) and (15) are not innate, but, rather, are responses to physical law, such that no group of subjects would respond differently, while example (11) is more naturally thought of as a learned behaviour. It is interesting to note that examples (9), (13) and (15) were the three examples with the highest level of agreement that the associated phenomena were not behaviours, therefore showing that our characterisation is also capturing an important element in our mental models.

Notes on the examples in table 3:

1. In this example there is no explicit environment or stimulus. The state change as response is to “not lay eggs”. The probability for this depends on several factors, as worker ants can lay eggs in certain circumstances [5]. However, taking queen ants as a benchmark, there is a clear distinction between the probability to lay eggs as a worker versus a queen. If we consider our ensemble to be both workers and queens, then with respect to this benchmark it is a behaviour, as there are multiple response to the same, implicit, stimulus. We can also see it as innate behaviour that is an adaptation that increases the inclusive fitness of the nest.
2. In this example there is no explicit environment, though we can imagine food as part of the environment. The considered response is pumping water. In the question of a benchmark we could take away the food stimulus. However, we would find that the sponge still pumped water, as it also does so for respiration and waste removal. i.e., there are multiple stimuli that lead to the same response. In this sense it is a behaviour. However, the probability to pump water is the same, independently of whether there is food or not. So, with respect to this benchmark, according to our characterisation, it is not a behaviour. It could be considered an innate behaviour with respect to some other benchmark, such as some other species that pumps water, but not do so with respect to the same food stimulus. For example, starfish, pump water through their vascular system in order to be able to move. As it is a very ancient adaptation, to view it as a behaviour from an evolutionary context one would have to determine those ancestors of sponges that did not pump water.
3. In the spider example, there is no explicit stimulus. The response is to build a web and this changes the environment from ‘no web” to ”web”. A natural benchmark is that of non-web-building spiders which allows us to identify it as a behaviour, as there are multiple responses in an ensemble of web-building and non-web-building spiders. The behaviour is innate and arises as an adaptation for prey capture.
4. There is no explicit environment and the stimulus is winter. A natural benchmark is that of rabbit species as subjects that live in more temperate or semitropical environments, where there is no need to grow thicker fur. In this case growing thicker fur is an innate behavioural adaptation that distinguishes different rabbit species. Of course, growing thicker fur would naturally be a behaviour when constructing an evolutionary ensemble that contained both those ancestors of rabbits before the adaptation occurred and their descendants where the adaptation was present.
5. For the plant stomata, there is no explicit environment or explicit stimulus. The response is to close pores. In spite of the lack of an explicit environment and stimulus, in this example an objective is added: ‘to conserve water”. It is an innate response and therefore an appropriate benchmark is one that takes into account its evolutionary origins. As it in an ancient adaptation [6, 7], designed to promote photosynthesis as well as conserve water, a relevant evolutionary benchmark would be pre-terrestrial ancestors of land plants. With respect to this benchmark it is a behaviour. However, if the subject and corresponding benchmark are both species of land plant, then with respect to this benchmark it would not be a behaviour. Given that stomata close at night, we could imagine choosing as benchmark the absence of the implicit stimulus “night” and consider what happens during the day. However, we would find that plants had the same change of response in the presence or absence of the stimulus and therefore could not use this to classify it as a behaviour.
6. For plant leaves bending towards light, there is no explicit environment and the stimulus is the presence of light. The response is to bend leaves towards the stimulus. In the case of phototropism, this is a very ancient, innate response, which is an evolutionary adaptation to maximise light capture for photosynthesis. A relevant evolutionary ensemble in which we could consider it as a behaviour would require considering ancestors of leafy plants before phototropism appeared. We could choose as benchmark the absence of the light stimulus, but this makes little sense, as the response involves the presence of the stimulus in the first place. However, an interesting benchmark with respect to which we can view leaves bending towards light as a behaviour is that of photonasty [8], which is the non-directional opening or closing of plant leaves and flowers in response to light intensity, rather than the direction of the light source.
7. The stimulus is a nightmare and the response is the heart beating faster. There is no explicit environment. Nightmares occur during REM sleep while night terrors occur during non-REM sleep. However, both can lead to the heart beating faster and harder. Thus, if we take as benchmark persons experiencing night terrors we would consider this example as a behaviour as there are multiple stimuli that lead to the same response.
8. Here the subject is a person, the stimulus is hot air and the response is to sweat. If we take as response a binary variable - sweat = yes/no and as benchmark a group of humans, then as all humans sweat this would not be a behaviour. This can also be seen by taking away the stimulus, say cold air, and also noting the uniform response across humans. From an evolutionary perspective however, sweating is an adaptation that has allowed humans to keep core temperatures at a tolerable level when subsistence hunting [9] and we can compare with benchmarks associated with other primates. Apes and monkeys sweat but lemurs do not. We can consider as evolutionary ensemble a last common ancestor that did not sweat and then descendants that did with the corresponding evolutionary advantage. If instead of considering as response sweat = yes/no we consider the amount of sweat, associated with the density of sweat glands, then there are clear differences with other primates as humans have a much higher density. In this case, taking as variable “sweating a lot” and comparing humans with other primates as benchmark subjects then this would indeed be a behaviour. It is known that, although the density of sweat glands is very high at birth, if the person is not in a hot climate, many of the glands become permanently inactive. Additionally, accumulated exposure to heat or exercise can cause more sweat glands to become active [10]. In this case we can take as subjects people in hot climates and those in cool climates as benchmark and notice significant differences in sweating. In this context sweating is clearly a behaviour.
9. In this example, the subject is a beetle, the stimulus a strong current and the response is to be swept away. Here, there is no relevant benchmark where there is an option to being swept away. Everything will be “swept away” with a sufficiently strong current. So, there is no other response possible and this is therefore not a behaviour.
10. The subject is a rat, the stimulus is salty food and the response is to dislike the food. This is not an innate response as if the rats are salt depleted then they will have a preference for the salty food. So, if we compare non-salt-depleted rats with salt-depleted rats as benchmark we will find different responses to the same stimulus and therefore this is a behaviour according to our characterisation.
11. Here, the subject is a person, the stimulus is the prediction of rain tomorrow and the response is to decide to not do anything. This is clearly not an innate response. In this case we could consider a longitudinal ensemble associated with the same person to determine if sometimes the person will decide to do something rather than nothing in spite of the rain prediction. Alternatively, we can consider a transverse ensemble, where we count how many people do something, even though rain has been predicted. In either case, it is clear we will find, though according to our prescription the corresponding ensemble should be created and measured, that there are multiple responses to the same stimulus and therefore this is a behaviour.
12. The subject is a horse, the “stimulus” is aging and the response is arthritis. Interestingly, this example had only 6% support for being a behaviour. This is understandable as in our mental models the horse has little choice in the matter. Clearly, all horses eventually with show signs of arthritis. Thus, using as benchmark one group of horses to be compared with another this will not be considered a behaviour. From an evolutionary perspective neither were there ancestors of horses that were not arthritic. In other words this is not an adaptation. However, there are various very long-lived vertebrate species, such as the Bowhead whale, Aldabra giant tortoise, Greenland shark and Rockfish, where the reported incidence of arthritis is much lower, with potential protecting factors being: very slow metabolism and low weight bearing stress in the case of aquatic vertebrates. In this case using any of these long-lived animals as benchmark would lead us to characterise this as a behaviour, albeit a somewhat unintuitive one as it is clearly not linked to a positive adaptation.
13. Here, the subjects are mice, the environment is outer space, the stimulus is zero gravity and the response is to float. In this case, there is no other group of subjects that would do other than float in that environment. The universal update rule here is Newton’s law. Hence, there is only one possible response to the stimulus and this is not an adaptation and the response is neither innate nor learned.
14. In this case, unicellular algae are the subject, food is the stimulus and the response is swim towards the nutrients. This is an innate response and a positive adaptation. If we consider as benchmark the absence of the stimulus, then we would see a difference in response.
15. Frogs are the subject, earth is the environment, gravity the stimulus and the response is to orbit the Sun along with the rest of the Earth. As with the floating mouse example, this is not a behaviour as there is no other response possible for any reasonable benchmark group of subjects.The universal update rule is Newton’s law and there is only one possible response to the stimulus as the response is neither innate nor learned.
16. The subjects are flocks of geese. There is no explicit environment or stimulus. The response is flying in 𝒱 formation. Two natural benchmarks here are non-migratory birds or migratory birds that migrate alone, such as cuckoos. The latter is a better benchmark if we add in the information that the birds are migratory. Thus, in an ensemble of geese and these benchmark birds it is clear that there is more than one response behaviour, for example to the stimulus that causes the birds to migrate, and, hence, this is a behaviour. It is an innate behaviour in the geese and is a positive adaptation to conserve energy during long migrations [11].
17. In this example, the subject is a dog, there is no explicit environment, the stimulus is anticipation of feeding time and the response is to salivate. Having the stimulus as anticipation of feeding time implicitly suggests that the feeding time has been set at the same time. In this case, there are at least two interesting benchmarks. Firstly, by changing the feeding time we will see that the response changes as the dog will be unable to anticipate, until it has learned the new feeding time. Hence, there are multiple responses to the same stimulus - feeding - and in this case it is a learned behaviour. Secondly, the dog salivates upon sensing food and this is an ancient, innate response that is a positive adaptation, as it aids digestion. In this case the appropriate benchmark is an evolutionary one, whereby we identify the evolutionary epoch before salivary excretion was used to aid digestion and compare it to one afterwards, with the combination being our ensemble. In this case *P* (*C*|*S, E*), where *C* is salivation, will change from ∼ 0 to ∼ 1.
18. The subject is herds of zebra, there is no explicit environment, the stimulus is breeding season and the response is breaking up for the season and reforming after it. This response depends on the zebra species, as mountain and plains zebras, don’t ”break up” for breeding season. Considering as subject zebra species that do break up and using as benchmark ones that don’t then there are multiple responses to the same stimulus and therefore this is a behaviour.
19. In this case the subject is a chameleon, the stimulus is sunlight and the response is to change colour. Evolutionarily speaking colour change is an adaptation that serves many purposes, including intraspecies signalling of emotions, as well as camouflage in some species. Although, all chameleons exhibit some degree of colour change in sunlight, the amount and type is highly variable depending on the species and its environment. As different stimuli can lead to the same response this is a behaviour.
20. Here, a cat is the subject, excess sugar in the blood the stimulus and insulin production the response. This is clearly an innate response associated with a positive and ancient evolutionary adaptation - to be able to use and store glucose. If we consider as benchmark an ensemble consisting of any other vertebrate species then this would not be a behaviour, as there is no multiple response to the same stimulus. As it is an ancient adaption present in all vertebrates a benchmark ensemble of invertebrates would have to be considered to be able to discern the possibility of multiple responses.

## Footnotes

1 The etymology of conduct is: mid-15c., “action of guiding or leading, guide”.

2 With this in mind, game theory would be one suitable framework in which to consider behaviour. In the case where we fix the space of rules and know what they are then this can be useful. However, in general we do not know much about the space of update rules other than what might be inferred from observing certain specific state changes in response to either observed or unobserved stimuli.

3 The reader may think that population genetics is a counter-example of these assertions. However, population genetics is such that for a given update rule, based on, say, selection, mutation and recombination, the response in gene frequency space is uniquely determined. In that sense, the dynamics associated with standard population genetics is more akin to applications of Newton’s Laws.

4 Note that various data sets are publicly available at the CHILAM website, as is a machine learning platform (project42.c3.unam.mx) that uses the Bayesian framework presented in this paper.

